# Distinct GSDMB protein isoforms and protease cleavage processes differentially control pyroptotic cell death and mitochondrial damage in cancer cells

**DOI:** 10.1101/2022.07.24.501218

**Authors:** Sara S Oltra, Laura Sin, Sara Colomo, María Pérez-López, Angela Molina-Crespo, Kyoung-Han Choi, Lidia Martinez, Saleta Morales, Cristina González-Paramos, Alba Orantes, Mario Soriano, Alberto Hernandez, Ana Lluch, Federico Rojo, Joan Albanell, Jae-Kyun Ko, David Sarrió, Gema Moreno-Bueno

## Abstract

The formation of Gasdermin (GSDM) pores, leading to pyroptosis or other context-dependent consequences, is directly involved in multiple diseases. Gasdermin-B (GSDMB) plays complex and controversial roles in pathologies, with pyroptosis-dependent and independent functions. GSDMB is promising oncologic therapeutic target since it exhibits either antitumor function, when immunocyte-mediated Granzyme-A (GZMA) cleaves GSDMB releasing its cytotoxic N-terminal domain, or pro-tumoral activities (invasion, metastasis, and drug resistance). However, it is still unknown the precise regulatory mechanisms of GSDMB pyroptosis as well as the differential effects of the four translated GSDMB variants (GSDMB1-4, that differ in the alternative usage of exons 6-7) in this process. Here, we first prove that exon 6 translation (in the interdomain protein linker) is essential for pyroptosis, and therefore, GSDMB isoforms lacking this exon (GSDMB1-2) cannot provoke cancer cell death. Consistently, in large series of breast cancer samples GSDMB2 expression, and not of exon6-containing variants (GSDMB3-4), associates with unfavourable clinical-pathological parameters. Moreover, cellular, and biochemical analyses combined with confocal, live cell imaging, and electron microscopy studies, demonstrated that diverse GSDMB N-terminal constructs containing exon-6 induce mitochondrial damage (increased mitochondrial ROS, membrane potential loss and mitochondrial DNA release) together with pyroptotic membrane cell lysis. While exon-6 residues are not required for membrane or mitochondrial localization, we also identified other key residues for N-terminal domain cytotoxicity. Additionally, we demonstrated that all GSDMB variants share the cleavage sites for GZMA, Neutrophil Elastase (identified in this study) and caspases. Interestingly, whereas Neutrophil Elastase and caspases produce N-terminal fragments in all GSDMB isoforms with no pyroptotic activity, thus acting as a potential inhibitory mechanism, GZMA cleavage activates pyroptosis in an isoform-dependent way. Summarizing, our results have important implications for understanding the complex roles of GSDMB isoforms in cancer and other pathologies and for the future design of GSDMB-targeted therapies.

## Introduction

Pyroptosis is an inflammatory form of programmed cell death that involves cell swelling and lysis, which causes massive release of cellular contents and thereby triggers strong inflammation (1, 2). Pyroptosis, which is activated in response to diverse damaging signals, has been involved in multiple pathologies, including infectious, inflammatory diseases and cancer (3–5). Pyroptotic cell death is also called “Gasdermin-mediated programmed necrosis” (6, 7), since the Gasdermin (GSDM) family of pore-forming proteins plays a key role in this process. Except PJVK, all GSDMs members (GSDMA/B/C/D/E) share a similar protein 3D structure and, a common pro-cell death activation mechanism (8–11). Briefly, the N-terminal (NT) pore-forming domain is auto-inhibited by the C-terminal (CT) domain, and GSDM proteins remain inactive in the cytosol. After specific stimuli, the released NT binds to diverse lipids and form pores in the cell membrane, as well as in mitochondria and/or other organelles, subsequently leading to cell death (4, 12–15). For each GSDM, the active NT is released, in a biological context-dependent manner, by the cleavage of the linker inter-domain region via specific cell-intrinsic or extrinsic proteases. For instance, GSDMA-NT by Streptococcal pyrogenic exotoxin B (16), GSDMB through lymphocyte-derived Granzyme A (GZMA) (17, 18), GSDMC by caspase 6/8, GSDMD by pro-inflammatory caspases (casp1/4/5/11) (19–22), apoptotic caspase8 (23), neutrophil elastase (NE) (24), or Cathepsin G (25), and GSDME via caspase3 or GZMB (18, 26). Moreover, alternative cleavage events can inactivate the NT pore-forming activity of some GSDMs (10, 27–29).

Accumulating evidences showed that GSDMB plays complex biological roles since it can exhibit either cell-death dependent and independent functions in diverse pathological conditions such as infection by enterobacteria (30), asthma (31, 32), inflammatory bowel disease (29, 33–35), and cancer (15, 36, 37). GSDMB is expressed in diverse organs (mostly in gastrointestinal tract, respiratory system, lymphoid tissues (38)) and in multiple tumor types (including breast, gastric and bladder, among others) (37). In tumors, GSDMB promotes either pro-tumor or anti-tumor functions depending on the biological context (37). GSDMB is frequently co-amplified with HER2/Erbb2 oncogene in breast carcinomas (39), where GSDMB over-expression promotes tumorigenesis (40), invasion, metastasis, and resistance to therapy (39, 41). Paradoxically, GSDMB can also have an anti-tumour role if its pore-forming pyroptotic function is activated in cancer cells either with a GSDMB-targeted nanotherapy (41) or extrinsically through its cleavage by GZMA (17). In a context of immune activation, the release of GZMA by NK and T-cells can cleave GSDMB either at the K229 or K244 residues within cancer cells, inducing pyroptosis and the subsequent enhancement of the antitumoral immune response (17). In fact, GSDMB is a potential therapeutic target in cancer and other diseases, being the activation of its pyroptotic activity a promising approach for efficient tumor killing. However, to develop future GSDMB-targeted treatment approaches it is essential first to fully define the precise functional domains and the regulatory mechanisms of GSDMB pyroptotic activity, since there are controversial and contradictory results. Aside GZMA cleavage (17), Panganiban and collaborators (37) proposed that caspase 1 cleaved GSDMB (after D236 in the linker region) releasing a pyroptotic NT fragment, while Chen et al (38) showed that almost all caspases could cleave GSDMB within the NT domain producing a cell-death inactive protein. Moreover, Shi et al (42) reported that GSDMB NT domain did not induce cell death, whereas Ding et al (8) generated a GSDMB construct of 275 aa (including part of the CT region) that produced strong lytic death, though whether any protease could generate this fragment was not demonstrated.

Of note, it is important to highlight that there are different GSDMB protein variants but their potentially distinct functional relevance in physiology and disease is usually overlooked. Indeed, GSDMB gene produces at least six transcripts (NCBI Gene ID: 55876) that are translated into four different protein variants (termed GSDMB 1-4), being their cell/tissue-specific expression pattern (GTEx Portal)(43) controlled by genetic features (diverse SNPs (32, 44)) and other unknown regulatory mechanisms (15, 45, 46). The four translated isoforms differ only in the presence of exon 6 (13 aminoacids, aa) and 7 (9 aa): GSDMB1 lacks exon 6 (Δ6), GSDMB2 lacks exons 6 and 7 (Δ6-7), GSDMB3 contains both, and GSDMB4 lacks exon 7 (Δ7). The residues translated by these exons are located within the flexible inter-domain linker region of the protein (10), but their biological function remained unclear so far. While recent reports indicate that these variants could mediate different effects both in cancer (47) and inflammatory diseases (31), their precise regulatory mechanisms and their involvement in pyroptosis and other GSDMB functions is largely unknown.

To shed light into the mechanism of GSDMB cell death induction herein we determined the GSDMB regions essential for pyroptosis and subsequently we described a differential role of the distinct GSDMB isoforms in this process. Moreover, our data indicate that GSDMB-driven cell death is associated with a concomitant mitochondrial damage. Furthermore, we have also demonstrated that specific proteases can generate different pyroptotic active or inactive GSDMB fragments. Importantly our data also revealed that the expression of GSDMB pyroptotic-proficient and -deficient isoforms differentially correlate with clinic-pathological parameters in breast carcinomas. This pioneering study clarifies the mechanisms of GSDMB pyroptosis and highlight the distinct relevance of GSDMB isoforms in cancer biology.

## Materials/Subjects and Methods

Detailed information of the methods is provided in **Supplementary Information.**

### Human samples

Breast cancer samples are part of a multicenter, prospective, observational study (48) coordinated by GEICAM (Spanish Group for Breast Cancer Research) and the participation of 31 Spanish hospitals. The study protocol was approved by the Institutional Review Board and the Ethics Committee of Hospital Provincial de Castellón (Spain), according to the requirements of the Spanish regulations (GEICAM 2009-03; clinicaltrials.gov identifier: NCT01377363). Study procedures were carried out in accordance with the Declaration of Helsinki, as revised in 2008, and good clinical practice guidelines. Written informed consent was obtained from all patients before enrollment. In those tumors quantitative expression of GSDMB isoforms were performed (see **Supplementary Materials and Methods**).

### Cell biology methods

HEK293T, SKBR3, 23132/87 and THP1 cell lines were grown according to the standard conditions. Transient transfection of all constructs and corresponding empty vectors (detailed in **Supplementary Methods**) were performed using lipofectamine 2000 (Invitrogen) according to the manufacturers’ protocol. To evaluate GSDMB cytotoxicity and mitochondrial damage, Lactate Dehydrogenase (LDH) release tests, MitoSOX, TMRE and flow cytometry Caspase 3/7-SYTOX assays were performed according to manufacturers’ instructions. Mitochondrial DNA release was performed essentially as described before (49).

To evaluate proteolysis induced by neutrophil elastase (NE), cell lysates from transfected HEK293T cells were mixed at indicated concentrations of rhNE (Sigma), followed by incubation at 37°C for 1h. When indicated, BAY-678 inhibitor was included. GSDMB cleavage bands were evaluated by In-Gel Digestion and reverse phase-liquid chromatography RP-LC-MS/MS analysis. Co-culture of HEK293T or SKBR3 cells with NK-92 cell line was performed to analyze the cleavage of GSDMB by GZMA as described before (17). The cleavage products were visualized by Western blot (WB). Uncropped original WB are included as **Supplementary Information**. All antibodies used in the study are included in **Supplementary Table 2.**

### Microscopy methods

Localization of different GSDMB constructs was visualized by immunofluorescence in transient transfected cells (72h) fixed with 4% paraformaldehyde and stained with the primary antibodies (**Supplementary Table 2**) as described before (41). Confocal microscopy images were captured by LSM710 microscope (ZEISS, Oberkochen Germany, EU), and processed by Fiji software (Image J 1.52). Alternatively, live cell tracking imaging was performed with doxycycline inducible vectors expressing GFP-tagged GSDMB constructs (detailed in **Supplementary Methods**). To evaluate the mitochondrial morphology changes induced by GSDMB constructs, correlative light and electron microscopy (CLEM) procedures were performed in 23132/87 cells.

Cells were seeded in a permanox Lab-Tek chamber slide (Nalge Nunc International, Naperville, IL) and transfected with doxycycline inducible vectors. After 6h of transfection, cells were induced with Doxycycline at 200ng/ml, incubated with red MitoTracker™ Deep Red FM (ThermoFisher) for 30 min at 37°C and fixed in 3 % glutaraldehyde. The confocal images were acquired with a Leica TCS SP8 HyVolution II (Leica Microsystems, Wetzlar, Germany). After fluorescence capture, slides were processed for transmission electron microscopy analysis, FEI Tecnai Spirit BioTwin (ThermoFisher Scientific company, Oregon, USA). Pictures were taken using Radius software (Version 2.1) with a Xarosa digital camera (EMSIS GmbH, Münster, Germany).

### TCGA data analysis

GSDMB isoforms mRNA expression from breast cancer patients (N=1 093) was analysed using data from TCGA. Correlation of GSDMB isoform expression and overall survival was obtained using “survival” R package (R Bioconductor). P-values were calculated using long-rank test. Hazard rates were calculated by Cox proportional-hazards model. GSDMB isoforms expression was scored according to the mean expression. Results were considered significant when p-value < 0.05.

### Statistical analysis

Analysis of statistical significance for the indicated datasets was performed with ANOVA or t-test using GraphPad Prism 5 (GraphPad Software, La Jolla, CA). Results were considered significant when p-value < 0.05.

## Results

### Only the GSDMB isoforms containing exon 6 present cytotoxic activity

To assess the functional importance of the different regions of GSDMB protein on cell death induction, in particular those encoded by the alternative exons 6 and 7, and to identify the minimal GSDMB NT fragment with pyroptotic activity, seventeen myc/HA-tagged GSDMB constructs were used (**Figure 1A**). These constructs were named following the exon codifying number and the aminoacid sequence of the longest translated GSDMB variant (GSDMB3, 416 aminoacids), thus the four translated isoforms, differing in the presence of exons 6 and 7, were termed 1-416Δ6 (GSDMB1), 1-416Δ6,7 (GSDMB2), 1-416 (GSDMB3) and 1-416Δ7 (GSDMB4). We also cloned different NT constructs ending after the translation of either exon 5 (1-220; shared by all isoforms), exon 6 (1-233), exon 7 (1-242), as well as the 1-275 fragment (containing the linker and part of the CT region encoded by exon 9), previously described as pyroptotic (8), and its respective isoform variants (lacking exons 6 and/or 7). Moreover, during the cloning process of 1-275, two spontaneous mutations were originated, and these constructs (1-275^H51N^ and 1-275L212P) were also included in the study. We later generated the same point mutations within the 1-233 fragment (1-233^H51N^ and 1-233^L212P^). Additionally, within 1-416 we introduced the A340D point mutation, an aminoacid change that release the NT-CT autoinhibition and induces pyroptosis in other GSDMs (8). Finally, we cloned the CT fragment (92-416) that occurs when caspase-3 cleaves (after D91) in the GSDMB-NT region (10).

**Figure 1.**
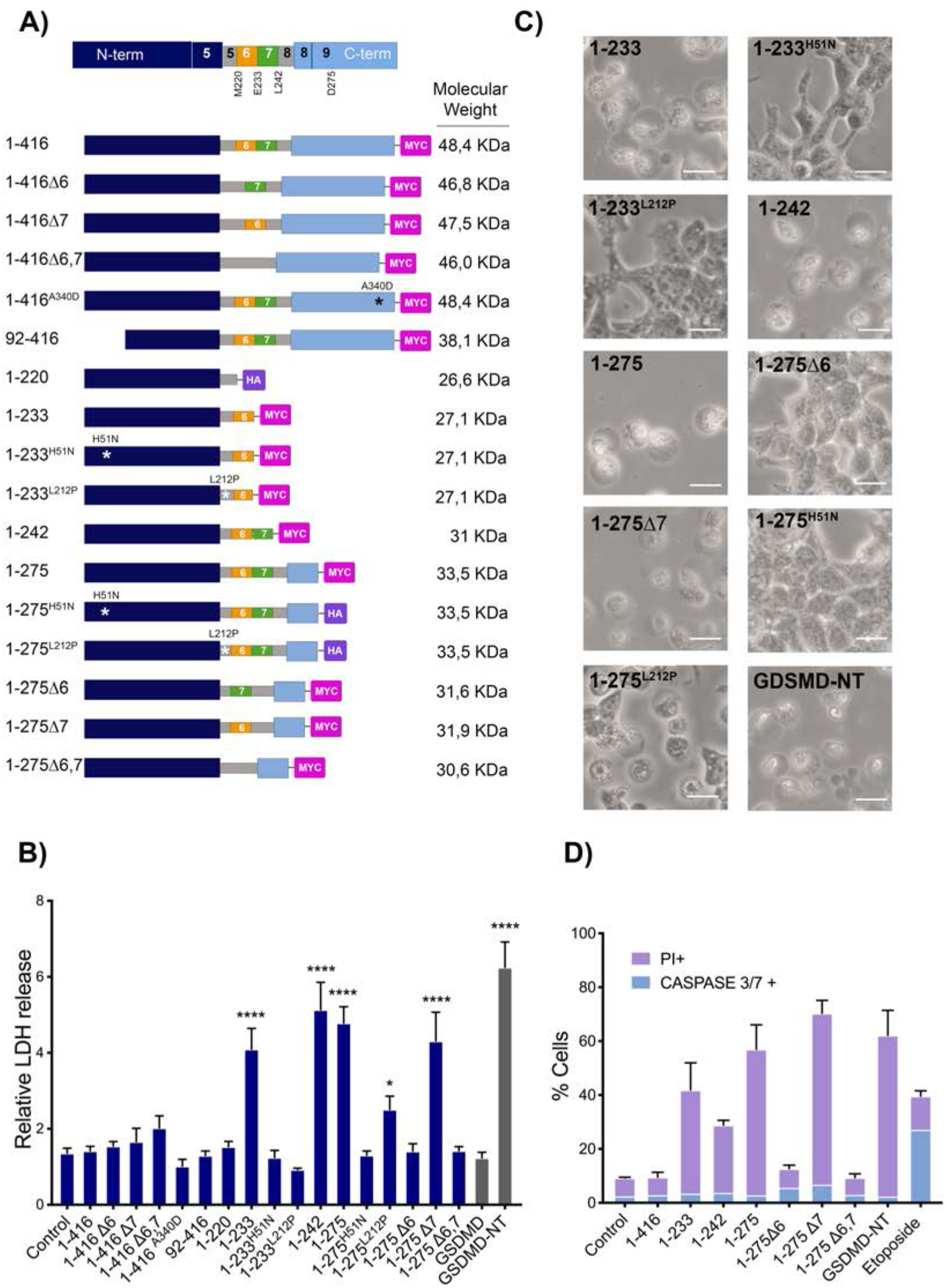
GSDMB exon 6 is essential for GSDMB-NT mediated pyroptosis. **A)** Top: Scheme of full length GSDMB protein, indicating the approximate location in the protein of the last residue encoded by exons 5 to 8. The flexible linker interdomain region (gray) comprises aminoacids Q210 to S252 and can contain the residues encoded by the alternative exons 6 (orange) and 7 (green). Below: schematic representation of the different GSDMB constructs used, indicating the localization of truncations or point mutations, and C-terminal tags. GSDMB translated variants GSDMB-1 to 4 are named 1-416Δ6, 1-416Δ6,7, 1-416, 1-416Δ7, respectively. **B**) Cytotoxicity of GSDMB constructs measured by lactate dehydrogenase (LDH) release assay in transiently transfected (48h) HEK293T cells. Empty vector (Control) and full-length GSDMD were used as a negative controls and GSDMD-NT as a positive control. Values are relative to empty vector and represent means ± SEMs of more than three independent experiments. Differences between control condition and each GSDMB construct were tested by t-test: *0,05 and ****p<0,0001. **C**) Representative bright field microscopy images of cells transiently transfected (48h) with the indicated GSDMB constructs. Scale bar represents 20μm. **D**) Apoptosis assays by flow cytometry analyses of caspase3/7 (light blue) activation (CellEvent™ Caspase 3/7 Green Flow Cytometry Assay Kit) and Sytox Green cell dye (purple). As positive control for apoptosis, cells were treated with etoposide (25μM) for 24h. Values represents the percentage of cells positive for each condition from at least three independent experiments.

To assess the pyroptotic potential of these constructs, they were transiently transfected (48h) in HEK293T cells, which do not express endogenous GSDMs (17) (**Supplementary Figure 1A and 1B**). Like the GSDMD-NT (1-275) pyroptotic domain (8), used as positive control, cells transfected with 1-233, 1-242, 1-275, 1-275^L212P^ or 1-275Δ7 constructs showed a significant increase in LDH release (**Figure 1B**) and exhibited pyroptotic morphology with remarkable cell swelling (**Figure 1C**). The remaining constructs presented LDH release levels similar to the empty vector controls and no pyroptosis morphology was appreciated (**Figure 1C**). Flow cytometry analyses of caspase3/7 activation and Sytox Green cell dye confirmed that GSDMB-NT cytotoxic fragments induced pyroptosis but not apoptosis (**Figure 1D)**.

These initial results provide novel insights into GSDMB pyroptotic function: **a**) Under unstimulated conditions, none of the full-length (not cleaved) GSDMB isoforms or a CT construct (92-416) previously described (10) produces lytic cell death; **b**) Among the NT constructs tested, the 1-233 (including exon 6) is the minimum fragment with pyroptotic activity, as 1-220 (up to exon 5) is not cytotoxic; **c**) The translation of alternative exon 6, but not exon 7, is required for pyroptosis, since all cytotoxic constructs (1-233, 1-242, 1-275, 1-275^L212P^ and 1-275Δ7) contains exon 6. In fact, removal of exon 6 but not exon 7, blocks cell death (compare 1-275 with 1-275Δ6, 1-275Δ7 or 1-275Δ6,7). This implies that only GSDMB isoforms 3 and 4 and not 1 or 2 (which lack exon 6) could have activatable pyroptotic activity; **d**) For GSDMB-NT cytotoxicity, the Histidine 51 residue is essential (H51N mutation completely blocks cell lysis in 1-233 and 1-275 fragments), while Leucine-212, has moderate relevance (L212P mutation inhibits 1-233 cytotoxicity but only partially reduces 1-275 cell death promotion (**Figure 1B**, **Supplementary Table 1**). Multiple sequence alignment of all GSDMs revealed that the Lysine-212 is highly conserved among GSDM family members (**Supplementary Figure 1C**). Although the Histidine-51 is not conserved, all the residues from the rest of GSDMs at this position belong to the aromatic family, which share similar properties. **e**) Finally, contrary to other GSDMs members (8), the mutation of a conserved Alanine-340 in the CT region (A340) does not release autoinhibition and activate pyroptosis of GSDMB.

### GSDMB NT-induced cell death associates with mitochondrial damage

Next, to assess the intracellular localization of our GSDMB constructs we performed immunofluorescence and confocal microscopy analysis using transiently transfected HEK293T cells. The 1-233 and 1-275 constructs with pyroptosis-inhibition mutations (1-233^H51N^, 1-233^L212P^, 1-275^H51N^ and 1-275^L212P^), as well as the 1-220 NT fragment localized in the cytosol but mostly accumulate as dot-like or ring-shape aggregates (**Figure 2A and Supplementary Figure 1D**). These aggregates mostly co-localized with the specific mitochondrial markers TOM20 (**Figure 2A**), HSP75/Trap-1 and mitoTracker Deep Red (**Supplementary Figure 2A**), and not with lysosomes (LAMP1) or the Golgi apparatus (GM130) (**Supplementary Figure 3**). Despite we were unable to visualize the most cytotoxic NT forms (1-233, 1-242, 1-275 and 1-275Δ7) by immunofluorescence in fixed cells, as dead cells readily detached from the coverslip, we confirmed their mitochondrial localization by western blot in purified mitochondrial fractions (**Figure 2B**). The remaining non-cytotoxic constructs were diffusely localized in the cytosol and showed no evident enrichment in any of the organelles tested (**Supplementary Figure 2 and 3**). Interestingly, the accumulation of GSDMB-NT fragments mainly associated with small mitochondria or with annular morphology (**Figure 2A** asterisk), suggesting a potential dysfunction of this organelle.

**Figure 2.**
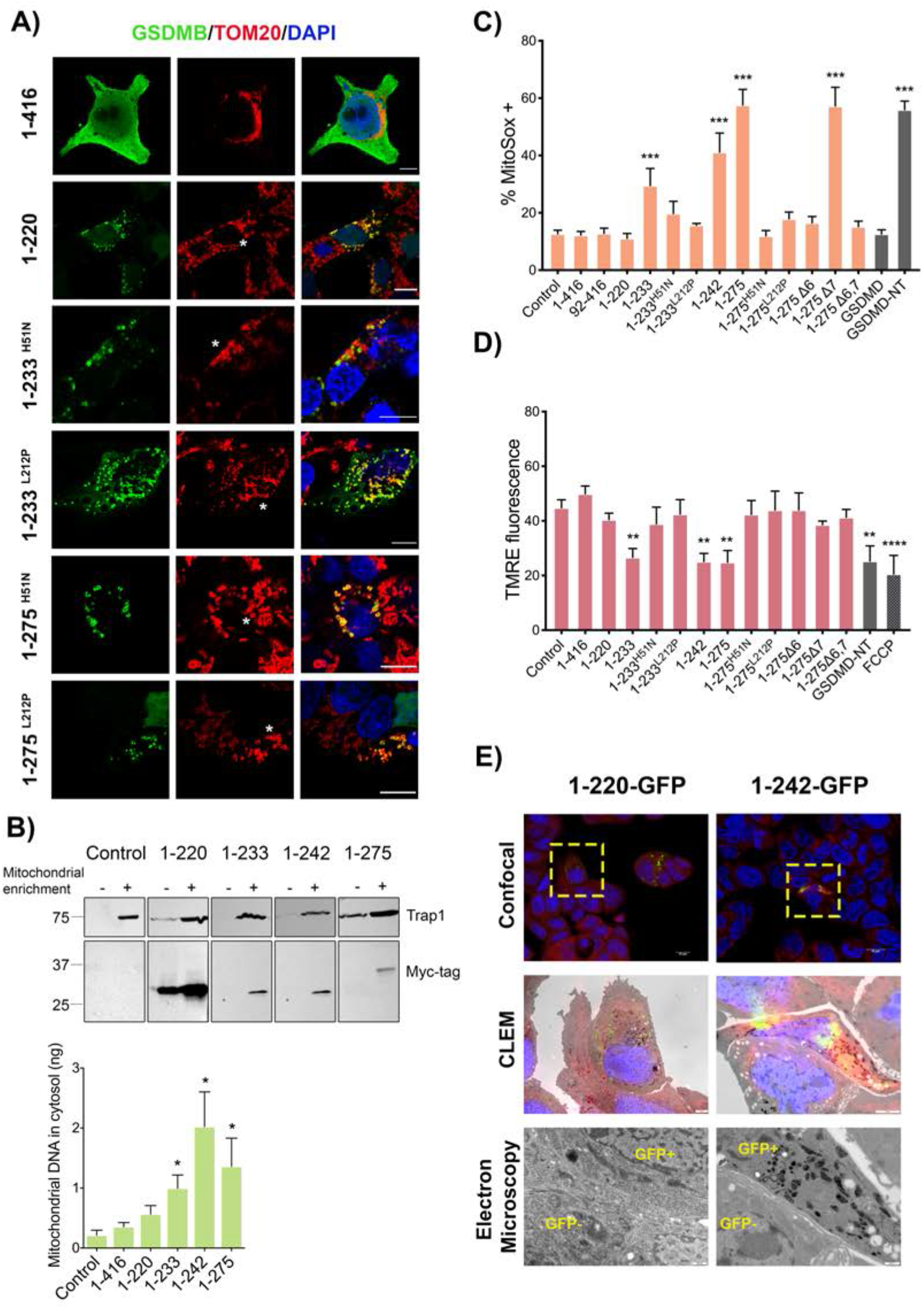
Mitochondrial localization of cytotoxic GSDMB-NT constructs associates with mitochondrial damage. HEK293T cells were transiently transfected (48h) with the indicated GSDMB constructs. **A)** Intracellular localization of GSDMB constructs (green; NT antibody SIGMA) and co-localization with mitochondria (red; TOM20) by immunofluorescence and confocal microscopy analysis. Note that GSDMB-NT constructs mostly localize in dot-like or round-shaped mitochondria (asterisks). Nuclei was stained with DAPI (blue). Scale bar represents 10μm. **B**) Mitochondrial localization of cytotoxic GSDMB-NT constructs by immunoblot in cytosolic and mitochondrial enriched fractions and quantification of mitochondrial DNA (mtDNA) presence in cytosolic fraction of HEK293T cells. Cytosol and mitochondrial fractions were analyzed by immunoblot probed for Myc-tag and Trap1. Mitochondrial damage was analyzed in HEK293T cells transiently transfected with GSDMB constructions during 48h for mitoSOX assay using flow cytometry (**C**) and during 24 h for Membrane potential TMRE assay by fluorescence (**D**). Values represent means ± SEMs of more than three independent experiments. Full length GSDMB construction, 1-416, was used as control because does not present colocalization with mitochondria. Differences between control and each GSDMB construction was tested by t-test: *p<0,05, **p<0,01 and ***p<0,001. GSDMD-NT was used as a positive control and, in TMRE assay, FCCP (carbonilcyanide p-triflouromethoxyphenylhydrazone) was used as a positive control for reducing mitochondrial membrane potential. **E**) Correlative light-electron microscopy (CLEM) study in 23132/87 cell lines. GSDMB-NT constructions 1-220-GFP or 1-242-GFP were transfected and induced with Doxycycline. Before CLEM study cells were targeted with mitoTracker Deep Red. Interest regions were selected (yellow square) by confocal microscopy and then processed for electron microscopy visualization.

These data are consistent with previous results in other GSDMs (GSDMA, GSDMA3, GSDMD, and GSDME) showing that their NT domains not only localize to the cell membrane (8, 22, 50, 51) but also target mitochondria (12, 14, 19), and likely other organelles (14). In fact, mitochondrial accumulation of GSDM NTs can induce diverse damage types such as rounding and fragmentation of mitochondria, mitochondrial DNA (mtDNA) release (49), membrane permeabilization, loss of membrane potential, and ROS (12, 42, 52–56). Therefore, to assess if GSDMB cytotoxicity also associated with mitochondria dysfunction we measured mitochondrial ROS (mitoSOX reagent), membrane potential (TMRE), and mtDNA release. Our data overall revealed a strong association between the cytotoxic capacity (LDH release) and the mitochondrial dysfunction in the constructs analyzed (**Supplementary Table 1**). The cytotoxic 1-233, 1-242, 1-275 and 1-275Δ7 fragments exhibited a sharp increase of mROS levels and decrease of membrane potential (except 1-275Δ7), like the GSDMD-NT positive control (**Figure 2C-D**, **Supplementary Table 1**). Interestingly, for the 1-233, 1-242, 1-275 cytotoxic fragments the ROS increasing occurs concomitantly with the mtDNA releasing at the cytosolic fraction, corroborating the mitochondrial damage enhancing (**Figure 2B**). Conversely, the corresponding pyroptosis-inactive mutants (1-233^H51N^, 1-233^L212P^, 1-275^H51N^) as well as the 1-220 fragment, produced no significant change in MitoSOX, TMRE levels or mtDNA release (**Figure 2C-D**), despite localizing at the mitochondria (**Figure 2A**). Unexpectedly, the 1-275^L212P^ fragment that exhibits moderate cytotoxicity (**Figure 1B**) showed no significant effect on MitoSOX and TMRE levels.

These alteration in ROS levels, mtDNA release and membrane potential could be related with mitochondrial structural modifications as was observed during apoptosis (57, 58). To analyze the mitochondrial ultra-structure, we performed the correlative light-electron microscopy (CLEM) study in 23132/87 cell lines expressing the GSDMB-NT GFP inducible constructions (1-220 or 1-242) that co-localized with mitoTracker Deep Red marker (**Figure 2E**). Compared to adjacent GFP-negative cells, the cells expressing 1-242 but not 1-220 exhibited higher electro-dense mitochondrial matrix. This feature has been related with an energetic functional form, producing energy by oxidative phosphorylation (**Figure 2E**) (59, 60).

Together, these results indicate that GSDMB-NT induced cell death associates with mitochondrial dysfunction. While highly cytotoxic fragments cannot be visualized by standard confocal imaging, point mutations inhibiting its killing activity allows its detection as aggregates into mitochondria. Despite abnormal appearance of mitochondria, these mutant constructs are incapable of producing measurable mitochondrial disfunction in terms of ROS and membrane potential. Moreover, the data proves that exon 6 is required for GSDMB-NT mediated cell death and mitochondrial damage, but not for mitochondrial localization, since the 1-220 construct (until exon 5) show the strongest mitochondria accumulation (and not mitochondrial damage, **Figure 2A-E**).

In addition to mitochondria targeting, our data prove that GSDMB-NT provoke cell lysis and release of intracellular LDH (**Figure 1**), suggesting that pores are formed in the cell membrane. Since we were unable to detect membrane localization by standard confocal immunofluorescence in transiently transfected and fixed cells, to track GSDMB intracellular localization and to determine the kinetics of pyroptosis and cell fate in real time, we performed time lapse fluorescence microscopy imaging with GSDMB constructs fused (in the CT) with green fluorescence protein (GFP) using doxycycline-inducible lentiviral vectors. For this, we generated the 1-416-GFP, 1-416Δ6-GFP, 1-220-GFP, 1-233-GFP, 1-242-GFP, 1-242Δ6-GFP, 1-275-GFP and 1-275Δ6-GFP constructs (**Supplementary Figure 4A-B**) and first proved that, after doxycycline induction, GFP-tagged proteins had the same effect that untagged ones in terms of cytotoxicity and mitochondrial ROS (**Figure 3A** and **3B**). The only exception was 1-233-GFP, which lost pyroptotic capability, likely due to GFP tagging, as reported before in a similar construct (19, 26). Besides, the new construct 1-242Δ6-GFP did not exhibit pyroptosis, as expected. Live cell imaging of HEK293T cells treated with doxycycline showed that (**Figures 3C and Supplementary Videos 1-12**): **a)** initially all constructs show diffuse localization, but only the NT fragments (not full-length 1-416, **Supplementary Videos 1 and 2**) gradually form aggregates (even ring-shape structures, mostly seen in 1-220-GFP, 1-275Δ6-GFP and 1-275 Δ6; **Supplementary Videos 3-8**) that increased in size with time; **b)** The aggregates move throughout the cell body and occasionally localize to the cell membranes (**Figure 3C**, **Supplementary Videos 3-8**); **c)** Importantly, membranous localization of 1-242-GFP and 1-275-GFP (arrows indicating) resulted in a visible membrane lysis, cell swelling and massive cell death as revealed by Propidium Iodide (PI) uptake (**Figure 3C** and **Supplementary Videos 9-12**) whereas augmented expression/aggregation or membrane localization of the other constructs did not provoke pyroptosis (**Supplementary Videos 3-8**). Enrichment in the cell membrane of diverse constructs was confirmed by Western blot (**Figure 3D**).

**Figure 3.**
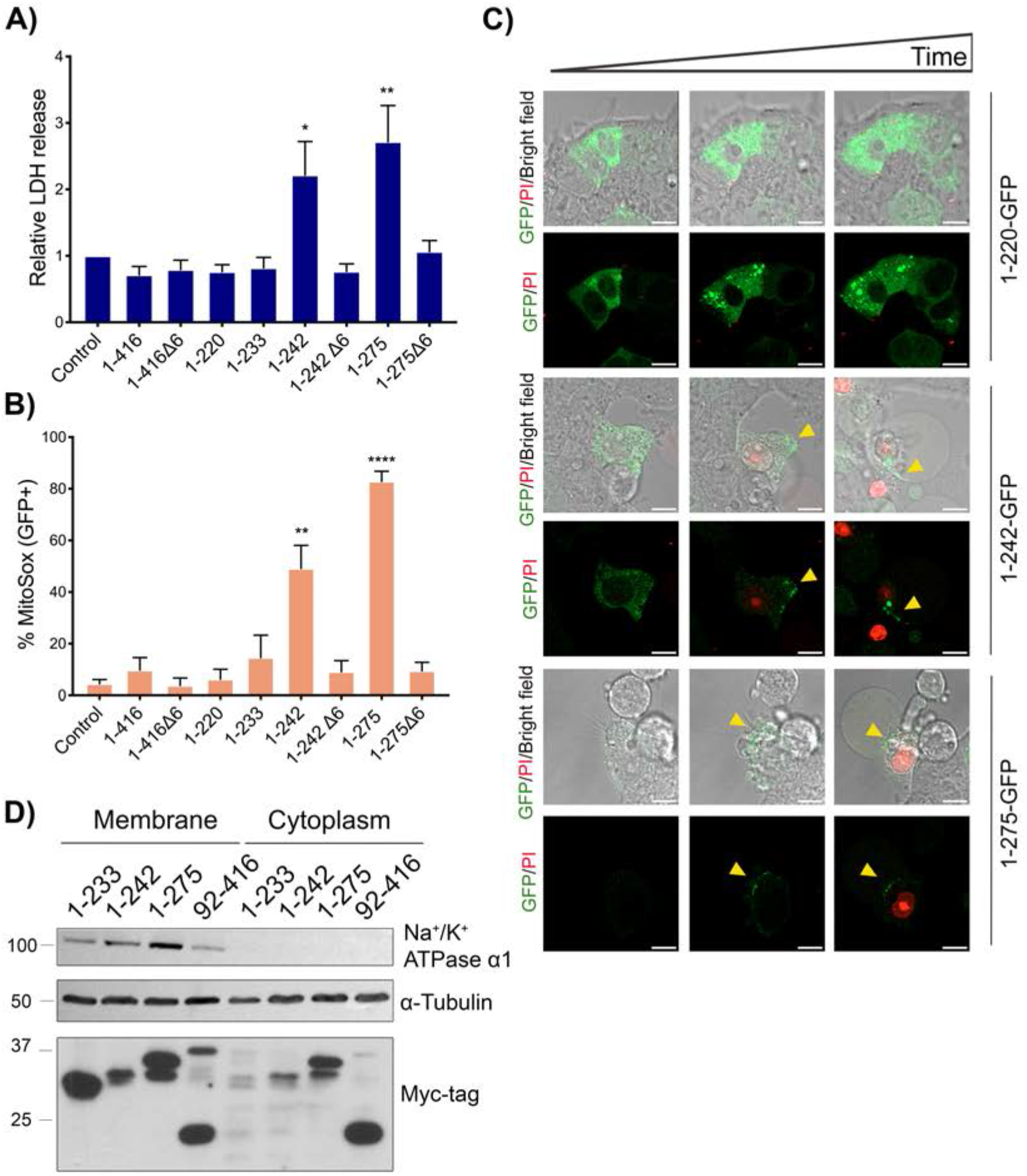
Cell membrane targeting of GSDMB-NT during pyroptosis. Doxycycline inducible GSDMB-GFP constructions were transiently transfected in HEK293T, and doxycycline induced after 24h of transfection. **A**) Cytotoxicity was measured using lactate dehydrogenase (LDH) assay after 16h of induction and **B**) Mitochondrial damage of GFP positive cells after 16h of induction was analyzed by mitoSOX using flow cytometry. Values represent means ± SEMs of more than three independent experiments. Differences between control condition and each GSDMB-GFP construction was tested by t-test: *p<0,05, **0,05>p<0,01 and ****p<0,0001. **C)** Microscopy imaging of GSDMB-NT localization in HEK293T during pyroptosis. Doxycycline inducible GSDMB 1-220-GFP, 1-242-GPF and 1-275-GFP were transiently transfected in HEK293T cells. Images are a time-lapse representation (brightfield and fluorescence) taken from 1-2 h after doxycycline addition. Scale bar represents 10μm. Green represents GSDMB and red Propidium Iodide. For complete videos see **Supplementary Videos 1-12**. **D**) GSDMB-NT cytotoxic constructions localize in plasma membrane. Membranes of HEK293T cells lysates, transfected as indicated, were enriched using Plasma Membrane Protein Extraction Kit (Abcam). Cytoplasm and membrane fractions were analyzed by immunoblot probed for Myc-tag, α-tubulin and Na^+^/K^+^ ATPase α1.

### GZMA, but not caspases, releases the GSDMB pyroptotic activity in an isoform dependent manner

Our data indicate that only exon 6-containing GSDMB isoforms (3 and 4) have activatable pyroptosis capability, but it is still unclear whether this function could be regulated (activate or inactivate) by differential protease cleavage.

To assess this question, we first focused on caspases since previous studies yielded conflicting results. Specifically, Panganiban et al (32) reported that caspase-1 could cleave after aminoacid D232, within exon 6, and release a pyroptotic GSDMB-NT specifically from the isoform 3, while Chen et al (29) showed that caspases 1/3/4/6/7/8/9 cleaved after residue D91, within the NT, producing two inactive GSDMB fragments. To clarify this issue, and since these studies utilized mostly forced co-expression of caspases and GSDMB in HEK293T cells, here we used THP1 monocytes stably over-expressing either GSDMB isoform 2 (1-416Δ6,7) or 3 (1-416). In these models, we induced canonical pyroptosis (standard LPS plus nigericin protocol (61)) or apoptosis (1μM etoposide for 24h). As shown in **Supplementary Figure 5 A-C**, our results proved that after activation of either pro-inflammatory (caspase-1) or pro-apoptotic caspases (caspase-3) GSDMB is cleaved in a region (D91 within NT) common to all isoforms. This caspase-dependent processing of GSDMB, which is analogous to the caspase-3 cleavage that inhibits GSDMD pyroptosis (27), produces GSDMB fragments with no effect on cell lysis (LDH release) during LPS+Nigericin induced pyroptosis (**Supplementary Figure 5 D)**. Thus, in agreement with Chen et al (29), and contrary to Panganiban et al (32), our data suggest that caspases do not activate GSDMB pyroptosis in an isoform dependent way.

Next, we studied GSDMB processing by GZMA, since a previous report showed that this protease could cleave two GSDMB Lysine residues, K229 (within exon 6) and K244 (exon 8). However, this study did not evaluate the differential functional effect on the pyroptotic activation of the GSDMB isoforms. To this end, HEK293 cells expressing each GSDMB isoform were co-cultured for 16h with NK-92 cells, following the reported method (17). Immunoblotting revealed an NT fragment of ~30kDa in all GSDMB isoforms, confirming the GZMA preference for the K244 cleavage site (23), common to all GSDMB isoforms (**Figure 4A**). Interestingly, we demonstrated that only exon 6-containing isoforms (GSDMB3 and 4) produced substantial increase of LDH release and mitoSOX levels in HEK293 co-cultured with NK cells (**Figure 4B-C**), thus demonstrating that GZMA-mediated pyroptosis induction depends on the GSDMB isoform and emphasizing again the essential role of exon 6 in this process.

**Figure 4.**
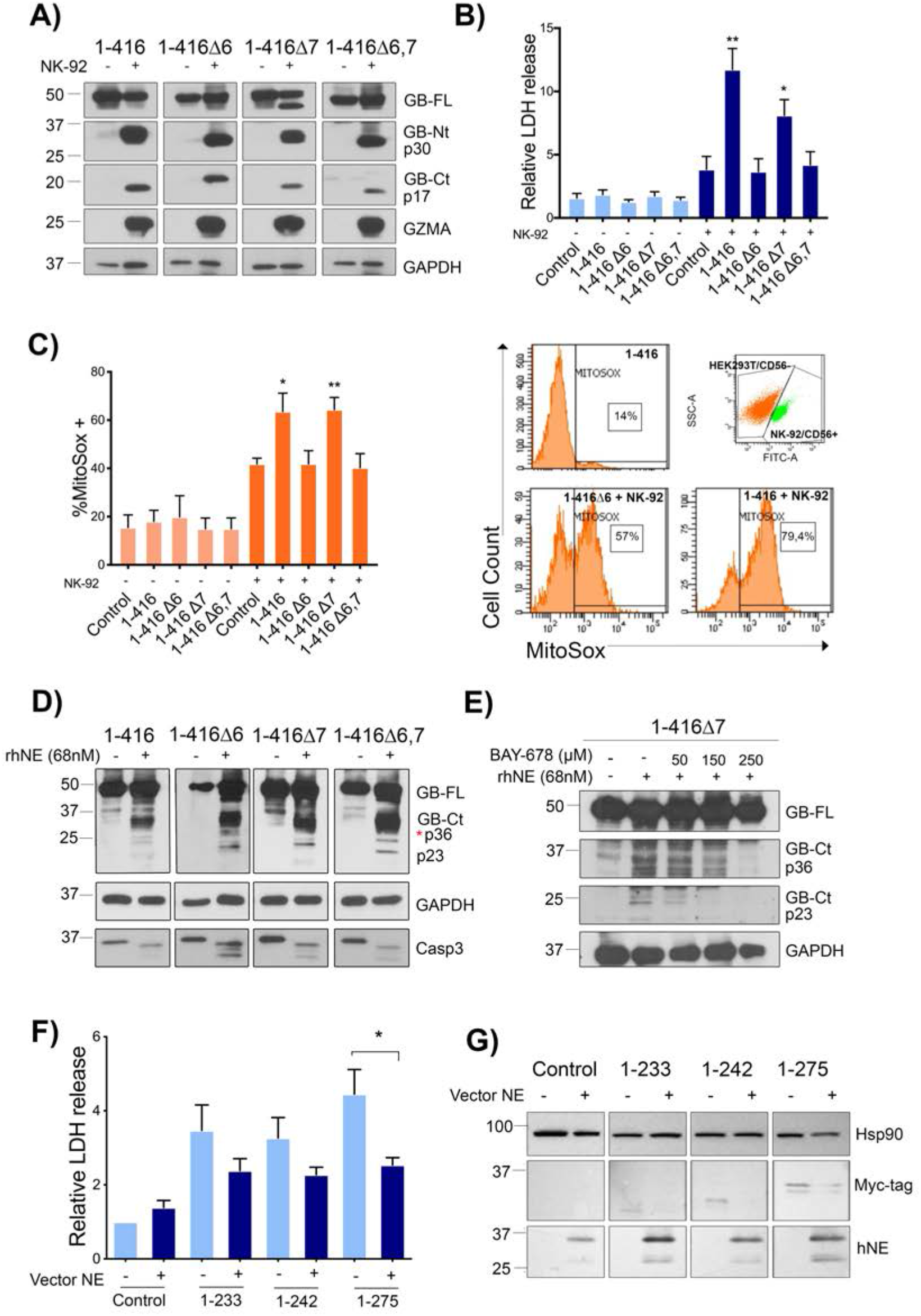
Cytotoxicity mediated by GSDMB-NT generated by processing by GZMA and NE cleavage depends on the exon 6 presence. **A)** Immunoblotting of HEK293T cells expressing GSDMB isoforms (1–4) and co-cultured with NK-92 cells to detect the cleavage of GSDMB by GZMA following established protocols (17) by immunoblot probed for anti-GSDMB-NT (Sigma, HPA023925) and anti-GSDMB-CT antibody DAN 114B (39) **B)** Lactate dehydrogenase (LDH) assay after 16h of NK-92 co-culture and **C)** Mitochondrial damage of HEK293T analyzed by mitoSOX using flow cytometry. NK-92 cells were labelled with CD56 antibody. Differences between control condition (HEK293T empty vector co-culture with NK-92) and each condition was tested by t-test: *p<0,05 and **p<0,01. **D**) Immunoblotting analysis of GSDMB cleavage by human neutrophil elastase (hNE). HEK-293T cell lysates containing GSDMB isoforms (1–4) were incubated with hNE (68nM) at 37°C for 30 min (**E**) and in presence of indicated amount of BAY-678 inhibitor. Immunoblot probes for GSDMB cleavage detection: anti-GSDMB-NT (Sigma, HPA023925) and anti-GSDMB-CT antibody DAN 114B (39). **F**) Lactate dehydrogenase (LDH) assay after 48h of co-transfection of hNE vector and GSDMB constructions. Differences between non-hNE and hNE presence for each condition was tested ANOVA multiple comparisons: *p<0,05. **G**) Co-transfection of HEK293T with hNE vector and subsequently with GSDMB constructions. GSDMB-NT constructions were cleavage by hNE producing a decrease in the GDSMB-NT fragments. Fragments were detected by anti-Myc flag located at the C-terminal region of GSDMB. All graph values represent means ± SEMs of more than three independent experiments.

### GSDMB cleavage by Neutrophil elastase blocks the pyroptotic activity

Finally, we studied the neutrophil elastase 2 (NE), a serin protease that cleaves and activates GSDMD NT pyroptosis in specific situations (24), since our preliminary *in-silico* analysis indicated that this protease could potentially cleave GSDMB.

Indeed, the *in vitro* digestion of protein lysates from GSDMB-expressing HEK239T cells with recombinant human neutrophile elastase (rhNE) resulted in the appearance of two major CT fragments (a large band ~36 KDa and a smaller one ~23KDa including the tag sequence) in all GSDMB isoforms (**Figure 4D**). Further information by mass spectrometry analyses revealed that the p23 fragment resulted from the cleavage at the residue M220 (exon 5 last amino acid), while the larger p36 corresponded to the CT fragment generated by caspases (10). Importantly, the addition of the NE specific inhibitor (BAY-678) in the *in vitro* reaction inhibited the generation of the p23 in a dose-dependent manner (**Figure 4E**), demonstrating that p23 cleavage was produced by rhNE, while p36 occurred from the spontaneous processing during *in vitro* incubation. Remarkably, cleavage by NE at M220 would produce the 1-220 NT fragment, which we have proved before to be non-cytotoxic (**Figure 1B**). Importantly, NE processing can inhibit GSDMB pyroptotic activity, since transient expression of NE (before transfection with GSDMB-NT fragments) reduced the cytotoxic effects of 1-233, 1-242, and 1-275 fragments in HEK293T cells (**Figure 4F**). Immunoblotting assay confirmed the reduction in the NT fragment, but unfortunately the cleavage fragments could not be detected (**Figure 4G**).

### GSDMB can kill cancer cells in an isoform-dependent way

Releasing GSDMB pyroptosis specifically in cancer cells can be a promising antitumor therapeutic approach. To assess this possibility, we first validated our results using two cancer cell models, 23132/87 (gastric cancer cell line) and SKBR3 (HER2 breast cancer) with very low or undetectable expression of GSDMB, respectively (**Supplementary Figure 1B**, SKBR3 cells harbor a *TATDN1-GSDMB* gene fusion, being the fused transcript predominantly expressed over the wildtype transcripts (62)). Replicating our results in HEK293T cells, 1-233, 1-242 and 1-275 constructs provoke pyroptotic cell death in SKBR3 (**Figure 5A)** and in 23132/87 cell lines (**Supplementary Figure 6A**), while the pyroptosis-inhibited mutant constructs accumulated in mitochondria in both cell lines (**Figure 5B** and **Supplementary Figure 6B**, respectively).

**Figure 5.**
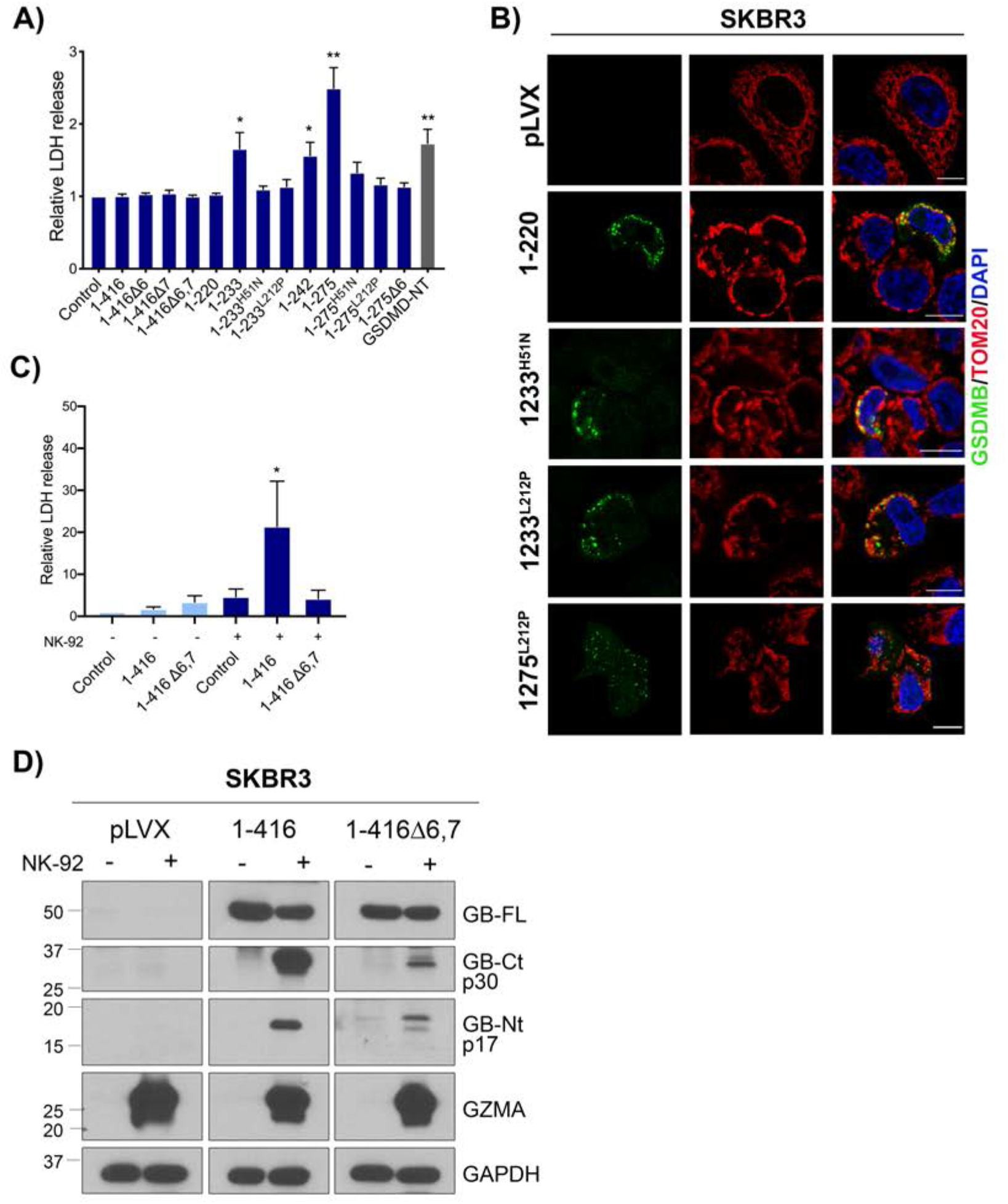
Pro-cell death role of GSDMB-NT in breast cancer cells in an isoform-dependent way. SKBR3 cell line was transiently transfected with GSDMB constructions during 48h. **A**) Cytotoxicity was measured by lactate dehydrogenase (LDH) assay. Differences between control condition (empty vector) and each condition was tested by t-test: *p<0,05 and **p<0,01. GSDMD-NT was used as a positive control. **B**) Immunofluorescence and confocal microscopy analysis in SKBR3 cells transiently transfected with indicated GSDMB-NT constructs. GSDMB-NT, indicated in green, colocalizes with mitochondrial marker TOM20 in red. pLVX was the empty vector used as a negative control. **C**) Lactate dehydrogenase (LDH) assay after 16h of coculture. SKBR3 cells expressing GSDMB isoforms 2 and 3 (1-416Δ6,7, 1-416, respectively) were co-cultured with NK-92 cells. Differences between control condition (SKBR3 empty vector coculture with NK-92) and each condition was tested by ANOVA multiple comparisons: *p<0,05. **D**) Immunoblotting of GSDMB cleavage by GZMA released by NK-92 cells co-culture with SKBR3 cells expressing GSDMB isoforms (2 and 3). Immunoblot probes for GSDMB cleavage detection: anti-GSDMB-NT (Sigma, HPA023925) and anti-GSDMB-CT antibody DAN 114B (39). All graph values represent means ± SEMs of more than three independent experiments.

Moreover, we also confirmed that NK-released GZMA can significantly increase cell death in SKBR3 in an isoform-dependent way (**Figure 5C-D**). Further corroborating the effect of NE cleavage in cancer pyroptosis, NE cleaves GSDMB in SKBR3 cells and releases the pyroptotic-inactive p23 NT fragment (**Supplementary Figure 6C**).

### Differential expression of GSDMB isoforms associates with clinicopathological variables in breast cancer

Based our results, we reasoned that the differential expression of GSDMB variants can have an impact on the biological and clinical behavior of GSDMB-positive tumors, since those mostly expressing exon 6-null isoforms (GSDMB1-2) should not have GSDMB-mediated pyroptotic anti-tumor function. To test this hypothesis, we focus on breast tumors, since we have previously demonstrated that in mammary carcinomas, GSDMB over-expression has prognostic value and promotes multiple pro-tumor effects *in vitro* and *in vivo* (39–41, 47). First, in 1 093 breast cancer patients from The Cancer Genome Atlas (TCGA) (63) we observed that GSDMB2 mean expression was generally higher than the other isoforms (**Figure 6A**). Consistent with this, GSDMB2 was significantly upregulated in breast cancer promoting tumorigenesis and metastasis *in-vivo* (47). Interestingly, in unselected breast carcinomas, GSDMB2 upregulation significantly associated with reduced overall survival, while higher levels of GSDMB1 and GSDMB4 (alternative expression of exon 6 and 7) have the opposite effect (**Figure 6B and 6C**). To validate these results, we assessed GSDMB2 isoform and GSDMB4 (with exon 6, as control) expression by qRT-PCR in 55 paired primary tumor and metastasis samples from the CONVERTHER clinical trial (NCT01377363), and again we observed that GSDMB2 expression in the metastatic lesions associated with significantly poorer overall survival. Although not significance was found in the GSDMB4 analysis, tumors with GSDMB4 high expression showed better outcome (**Figure 6B and 6D**)

**Figure 6.**
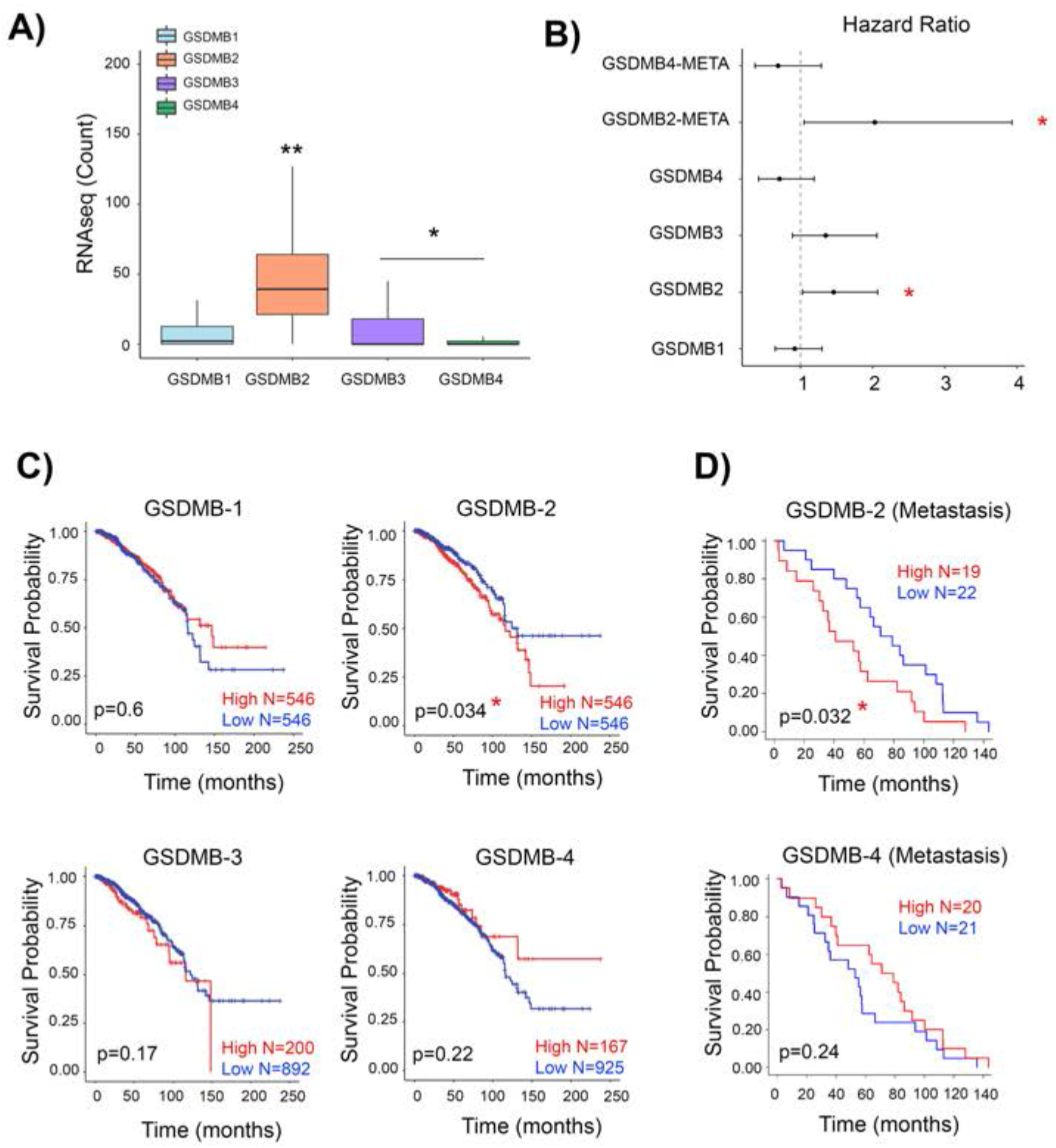
GSDMB-2 isoform expression associated with poor prognosis in breast cancer patients from TCGA and the clinical trial NCT01377363. **A)** mRNA expression of GSDMBs’ isoforms from breast cancer patients (N=1 093) was analyzed using TCGA data. GSDMB-2 isoform was significantly overexpressed in comparison with the rest of isoforms. Statistical analyses were performed by ANOVA and Tukey comparisons: *p<0,05 and **p<0,01. **B**) Hazard ratio representation. Hazard ratios were calculated by Cox proportional-hazards model. Survival probability graphs of GSDMs’ isoforms in all breast cancer patients from TCGA (**C**) and for GSDMB2 and GSDMB4 in metastatic patients from the clinical trial NCT01377363 which were analyzed by real-time PCR (**D**). Survival study was performed using “survival” R package. P-values were calculated using long-rank test. GSDMB isoforms expression was scored according to the mean expression. Results were considered significant when p-value < 0.05.

These results are in line with the enhanced *in vivo* aggressiveness and tumorigenic potential of GSDMB2 observed in breast cancer xenografts (47) and GSDMB2/HER2 knock-in mouse models (40).

## Discussion

GSDM-mediated pytoptosis participates in the genesis and progression of multiple diseases (4, 11). Therefore, the GSDMs are promising therapeutic targets with several GSDM-targeted approaches currently under pre-clinical evaluation (37, 64). Human GSDMB (mice and rat do not have *GSDMB* orthologue (65)), is involved in the response to enterobacteria infection (30), inflammatory diseases (asthma, inflammatory bowel disease or arthritis, among others; (29, 31, 32, 35, 66)) and cancer (15, 37). Surprisingly, in each of these pathologies contradictory GSDMB effects have been reported, and both GSDMB pore-forming-dependent (17, 30, 32) and independent functions has been described (29, 31, 35, 37). GSDMB plays a complex dual role in cancer, since GSDMB overexpression induce multiple pro-tumor activities in breast and bladder cancer, among others (15, 39, 41, 47, 67), but it can also exhibit a cytotoxic effect in a context of immune system activation (17).

To explain these conflicting results and to boost the chances of GSDMB-mediated therapies becoming a clinical reality in a near future for the treatment of cancer and other diseases, there are three key questions to be answered: a) differential effects of each translated GSDMB isoform (GSDMB1-4) in the diverse functions; b) mechanisms controlling pyroptotic and non-pyroptotic activities; c) protein regions and the intracellular events involved in each of these functions. The results described in this manuscript shed light into these important aspects.

First, in three different epithelial cell models (HEK293T cells, breast, and gastric cancer cell lines), we demonstrated for the first time that exon 6 (encoded by 13 amino acids) is essential for GSDMB mediated pyroptosis. This result has a strong relevance for understanding GSDMB functional involvement in multiple pathologies, since GSDMB pro-cell death activity depends on the translated isoform, being the variants lacking exon 6 (GSDMB1-2) pyroptotic-deficient. Thus, in cancer, the pro-tumor or antitumor activities of GSDMB may depend on the balance of their translated isoforms. We proved that expression of GSDMB1-2 would be beneficial for tumor cells, as these variants are resistant to pyroptosis-activation mediated by GZMA after immune attack. Indeed, the shortest variant (GSDMB2 which lacks exons 6-7) is particularly advantageous for breast cancer cells, as it increases *in vivo* tumor development and metastasis in MCF7 cell xenografts in immunodeficient mice and enhances tumorigenesis in an immune-proficient Knock-in model of GSDMB2/HER2 breast cancer (40). Consistent with this, GSDMB2 mRNA is the most upregulated transcript in the TCGA breast cancer cohort (63) significantly associates with unfavorable clinical-pathological and prognostic markers. Importantly, the poor prognosis value of GSDMB2 was confirmed in a breast cancer samples from a clinical trial (clinicaltrials.gov identifier: NCT01377363 (48)). Despite the limitations of mRNA studies (diverse variants can be expressed at the same time in a tumor, and unfortunately, there are currently no commercially available antibodies that could distinguish these isoforms by IHC in clinical specimens), we can speculate that while exon 6 is key for inducing tumor pyroptosis (and hence anti-tumor function) exon 7 might be important for other biological functions of GSDMB, such as transcriptional regulation. In this sense, the two exon 7-containing isoforms, GSDMB1 (in human bronchial cells) and GSDMB3 (in a GEMM hGSDMBZp3-Cre model), can mediate the transcriptional regulation of the same set of genes that are important for asthma (31).

The mechanisms that control the differential transcription, translation, and protein cleavage of GSDMB variants are largely unknown. At the transcription level, diverse SNPs can alter the balance of the isoforms (66), but these may have different effects in healthy and tumor tissue. In this sense, Lutkowska et al found that the rs8067378 G/G genotype upregulated GSDMB1 expression in both non-cancerous and cancerous cervical tissues but increased the other three isoforms only in the tumor (44). Of note, Panganiban et al reported that rs11078928-C allele (associated with asthma risk) produced exon 6 skipping and prevented the translation of a caspase-1 cleavage site at D236, thus producing pyroptosis-deficient GSDMB-NT in bronchial epithelial cells (32). Nonetheless, our data in THP-1 cells (over-expressing either GSDMB2 or GSDMB3), together with the results of previous studies (10, 29), prove that neither inflammasome-specific caspase-1 nor apoptotic caspases cleave D236 residue, but they cut D91, producing a short NT fragment with no pyroptotic activity. Aside SNPs, we investigated further other proteases that could differentially activate or inactivate GSDMB protein isoforms and confirmed that GZMA preferentially cleaved at K244 (common residue to all GSDMB variants) not the K229 (exon 6) (**Supplementary Figure 7A-B**). Thus, GZMA does not specifically cleave any isoform, but its activation effect on pyroptosis depends on the presence of exon 6-containing GSDMB variants. We also demonstrated for the first time that NE, which activates GSDMD in specific circumstances (24, 25, 68), can also cleave all GSDMB isoforms, in the M220 residue (translated by a shared codon between exon 5-6) (**Supplementary Figure 7A-B**), generating an NT domain with no cell-death effect. In fact, our data suggest that NE-cleavage could be a mechanism for defending epithelial cells and tumors from GSDMB pyroptosis by inhibiting the effect of GSDMB-NT cytotoxic fragments.

Regarding the precise GSDMB-NT region that induces pyroptosis, there are still conflicting data in the literature (8, 17). The minimum pyroptotic portion we identified was 1-233 peptide, which was as cytotoxic as 1-242 and 1-275 fragments (identified by (8)). As commented before, the killing effect of these fragments depends on the presence of exon 6 residues, despite these aminoacids lie within the linker region and not in the *bona fide* NT structural domain (1-225; (10)). In fact, the 1-220 construct, which does not include any aa of the flexible interdomain, is not cytotoxic. It should be noted that some discrepancies previously reported in the literature can be explained by the fact that adding tags to constructs around the minimal cytotoxic region can impede pyroptosis. Thus, Panganiban (32) reported that untagged 1-232 fragment was cytotoxic, while 1-233 construct fused with GFP ((19) and our data) or 1-236 tagged with N-terminal 3xFLAG (29) are not. This indicate that exon-6 might be important for maintaining a correct pyroptotic domain configuration. Additionally, our comprehensive study in 17 constructs also demonstrated that equivalent residues in GSDMB and GSDMD can have either similar or disparate effects on pyroptosis. Thus, H51 and L212 NT aminoacids are required for GSDMB cytotoxicity, and the corresponding/neighbor residues W48E, W50E and L192E in GSDMD, also prevented pore functionality (69). However, contrary to other GSDMs (8), mutating a conserved CT Alanine (A340 in GSDMB; A338 in GSDMA or A377 in GSDMD) to Aspartic does not release auto-inhibition and activate pyroptosis.

Moreover, our study also provides new information on the intracellular mechanisms of GSDMB cell death, as we proved that cytotoxic GSDMB-NTs induce two likely complementary processes, in one hand mitochondrial damage (including oxidative stress, loss of membrane potential, mitochondrial DNA release and electro-dense mitochondrial matrix) and in the other hand cell membrane rupture (and LDH release). This double effect is shared by other GSDMs (GSDMA/A3/D/E) but with some differences. Thus, contrary to GSDME-mitochondrial pore formation that provokes a positive feed-back loop by activating apoptotic caspase-3/7, GSDMB pyroptosis is not associated with caspase-3/7 secondary activation. Moreover, our confocal studies showed that diverse GSDMB-NT constructs form cytosolic aggregates that increase in size with time. Similarly, large cytosolic aggregates have also been reported for diverse GSDMD-GFP constructs (70) supporting that elimination/mutation of CT domain would result in insoluble GSDM-NT peptides. Importantly, these GSDMB-NT accumulations mostly co-localize with fragmented or rounded mitochondria, and electron microscopy confirmed that pyroptotic GSDMB NT fragments increased the mitochondrial matrix electro-density, which is related to an increased in the oxidative phosphorylation and subsequent generation of ATP (59, 71). This is likely a mechanism of cell recovery from the injury caused by the pyroptosis induction (60, 71). Furthermore, live cell imaging additionally proved that focal and transient localization in the cell membrane, specifically of exon 6-containing cytotoxic GSDMB-NT-GFP constructs, results in robust cell lysis, in a way akin to pyroptotic NT constructs of GSDMD (51) or GSDME (12). More precise single cell imaging studies of GSDMD pyroptosis dynamics showed that mitochondrial and lysosomal damage preceded cell lysis (14). Additionally, GSDMD can also target azurophilic granules and autophagolysosomes (72) or possibly also the nucleus (68), indicating that activated GSDMs could target diverse organelles under specific circumstances. Our study in pyroptotic-deficient point mutants show GSDMB-NTs mostly accumulate into mitochondria, and not lysosomes and Golgi, although is possible that GSDMB could also have context-dependent effect in other biological membranes, like pathogenic intracellular bacteria (30). The mechanisms that control GSDM-NT preference for mitochondria are currently unknown, but it might be in part mediated by their binding to cardiolipin (enriched in mitochondria) (8, 22, 73). However, the precise affinity of GSDMB to this lipid is controversial (10, 30), and contrary to other GSDMS, full length GSDMB could also mediate biological functions by binding to sulfatides (10, 41). Our data prove that 1-220 NT fragment is sufficient for mitochondrial localization but not for cell damage, while exon 6 region is required for mitochondrial damage and cell lysis but not for mitochondria targeting. However, the precise residues involved in lipid recognition and pore-formation are still unknown since GSDMB protein is not fully crystallized yet.

Summarizing, the data presented here together with other evidence prove the differential role of GSDMB isoforms in mediating diverse GSDMB biological activities. Whereas some pro-tumor functions might be common to all four isoforms, like cell motility (35, 41, 47) and possibly resistance to anti-HER2 therapies; (74) pyroptosis-mediated antitumor role is specific to GSDMB3-4.

Moreover, our novel data can help not only to understand the complex GSDMB roles in pathologies but to develop more precise *in vivo* models (including GEMM) for each disease, as well as to design future therapeutic approaches that could activate (in tumors) or inactivate (inflammatory diseases) GSDMB pyroptosis.

## Supporting information

Supplementary Materials and Methods, Supplementari figures and tables, supplementary references

## Acknowledgements

We are grateful to members of Gema Moreno-Bueno’s laboratories for constructive suggestions. We also thank Paloma Narros and Dr Javier Egea from UAM for helping us with the mitochondrial fractionation experiments. This study has been supported by the Ministerio de Ciencia, Innovación y Universidades, Agencia Estatal de Investigación (MICINN-AEI, PID2019-104644RB-I00) -GMB-, the Instituto de Salud Carlos III (CIBERONC, CB16/12/00295 -GMB- and -JA-, CB16/12/00241 and by the AECC Scientific Foundation (FCAECC PROYE19036MOR and GCTRA18014MATI -GMB-), the National Institutes of Health (NIH, USA) (R01-DK123475) -J-KK- and it has been also supported by a startup fund to -J-KK-from the Ohio State University, the College of Medicine, Department of Surgery. DS contract is funded by CIBERONC partly supported by FEDER funds. SO is funded by the FCAECC (POSTD20028OLTR), SC is funded by the MICINN-AEI PRE2020-095658. We are also thankful to all participating patients in CONVERTHER trial as well as the whole network of investigators and GEICAM staff for their contribution.

## Conflict of Interest

AL declares advisory boards and consulting fees from Novartis, Pfizer, Genentech, Roche, Eisai, Celgene; Research funding-clinical trials for institution from Novartis, Pfizer, Roche/Genentech, Eisai, Celgene, Roche Pharma AG, AstraZeneca, Merck, PharmaMar, Boehringer Ingelheim, Amgen, GlaxoSmithKline, Pierre Fabre. FR has received research grants from Roche; Consulting/advisory fees from BMS, MSD, AstraZeneca, Janssen, Roche/Genentech, Novartis, Eli Lilly, and Daiichi Sankyo. JA has received consulting or advisory role fees from Roche, Pfizer, Amgen, MSD, Lilly and Daiichi-Sankyo; research funding or grant support trials by Roche, Pfizer, Amgen, MSD, Lilly, Daiichi-Sankyo; and travel and accommodation support from Roche, Pfizer, Amgen, MSD, Lilly and Daiichi-Sankyo.

## Author Contribution

SO, DS and GMB conceived and designed the study. SO, LS, SC, MP-L performed most of the experiments, analyzed results and interpreted data. LS, SM, CGP participated in cell biology and biochemical experiments. AMC, K-H C and J-K K generated different GSDMB constructs and helped in data interpretation. AO and DS performed experiments with THP1 cells. MS and AH carried out the CLEM methodology. AL, FR, JA selected and followed up patients included at CONVERTHER clinical trial. SO, DS and GMB wrote the manuscript. All authors reviewed and edited the final manuscript.

