## Supplementary Materials and Methods, Supplementari figures and tables, supplementary references for "Distinct GSDMB protein isoforms and protease cleavage processes differentially control pyroptotic cell death and mitochondrial damage in cancer cells"

**Running title:** Different role of Gasdermin B isoforms in pyroptosis.

Sara S Oltra, Laura Sin, Sara Colomo, María Pérez-López, Angela Molina-Crespo, Kyoung-Han Choi, Lidia Martinez, Saleta Morales, Cristina González-Paramos, Alba Orantes, Mario Soriano, Alberto Hernandez, Ana Lluch, Federico Rojo, Joan Albanell, Jae-Kyun Ko, David Sarrió*, Gema Moreno- Bueno*

*Corresponding authors

**INDEX**


- Supplementary Materials/Subjects and Methods 3
- Supplementary Figures 11
- Supplementary Tables 22
- Supplementary Videos legends 24
- References 26

### **Supplementary Materials/Subjects and Methods:**

#### **Plasmids**

The pEZ-M61 plasmid with a HA flag at the C-terminal region was used to generate 1-220 and 1-275 constructs. Unfortunately, as previously described (1), 1-275 construct was highly toxic in *Escherichia coli* and could not be cloned in the pEZ-M61 plasmid that presented a bacterial T7 promoter. Furthermore, during the cloning process spontaneous mutations of the 1-275 construct arose at c.153 C>A (p.His 51Asn) located in exon 2, and at c.637 G>A (p.Leu212Pro), located in exon 5, named as H51N and L212P, respectively. The rest of cDNA constructs were generated using the lentiviral vector pLVX with a C-terminal *myc* flag, a cytomegalovirus promotor and puromycin resistance. The cDNA was cloned in the lentiviral vector pLVX following the Ligase T4 DNA protocol (Invitogen). The pEZ-M61 vector was obtained from Genecopoeia (Rockville, MD, US), constructpLVX from Takara (Takara Bio, USA) and Neutrophile Elastase cloned in the pCMV6-XL4 from Origene (Rockville, MD, US). Inducible Lentiviral Vector tagged with GFP at the C-terminal was used for the inducible GSDMB fragments (Inducible Lenti-TRE3G-ORF-C-TagGFP2-PGK-Tet3G-puro; from Transomic) (**Supplementary Table 2**).

Transformation was performed in DH5α bacteria following the conventional thermal shock transformation protocol. cDNA was isolated by High Pure Plasmid Isolation Kit (Roche) and all plasmids were verified by DNA sequencing. As additional controls, GSDMD full-length cDNA (pCS2-3XFlag-hGSDMD) and GSDMD-NT (pCS2-3XFlag-hGSDMD 1-275) were kindly provided by Dr. Pablo Pelegrin (Instituto Murciano de Investigación Biosanitaria Virgen de la Arrixaca, IMIB, Murcia, Spain).

#### **Cell culture and transient transfection**

HEK293T cells were grown in Dulbecco's modified Eagle's medium (DMEM) supplemented with 10% fetal bovine serum (FBS) and 2 mM L- glutamine. SKBR3 and 23132/87 cell lines were grown in RPMI medium supplemented with 10% FBS and 2 mM L- glutamine. NK-92 cells were growth in 75% alpha-MEM supplemented with 12.5% FBS, 12.5% of horse serum, 2 mM of L-glutamine and 5ng/ml of IL-2. THP1 cells were cultured in RPMI medium supplemented with 10% FBS, 1% streptomycin/penicillin, 1% glutamine and 0.5% fungizone. All cells were grown at 37°C and 5% of $CO_2$. For co-culture assays, target HEK293T or SKBR3 cells were seeded in 12-well plates (3 x $10^5$ cell/well) and transfected for 48h. Then, NK-92 cells were added at 8:1 ratio (Effector cell: Target cell) and co-cultured for 16h. Cell lines were purchased from commercial repositories (Suppl table 2) and their identity was confirmed by short tandem repeat profiling. Cells were routinely tested for Mycoplasma infection.

Transient transfection of all constructs and corresponding empty vectors were performed using lipofectamine 2000 (Invitrogen) according to the manufacturers' protocol. For those cells transfected with the constructs cloned into Doxycycline-inducible lentiviral vectors, we added Doxycycline at 200ng/ml to induce gene expression at intended time frames. Regarding the neutrophile elastase assay, cells were firstly transfected with NE and 24h later with GSDMB constructs.

**Lactate dehydrogenase (LDH) cytotoxicity assay**

LDH release was assayed using the Cytotoxicity Detection KitPLUS LDH (Roche) following the manufacturer's protocol. Culture media from transfected cells, as indicated previously, was collected and centrifugated at 1 200 rpm, 4°C for 5 min to remove cell debris. LDH release was measured at OD 490. Relative LDH release was expressed as LDH release in transfected cells relative to LDH release in control cells (not transfected).

**MitoSOX and TMRE assays**

MitoSOX Deep red fluorescent probe (Thermo Fisher Scientific™, Carlsbad, CA, USA) was used to determine the superoxide production by the GSDMB constructs in cell lines. Briefly, superoxide levels were measured by mitoSOX in transiently transfected cells (48 h). After removal of transfection medium cells were incubated in 5µM mitoSOX for 30 min at 37°C. The mitoSOX levels were measured by flow cytometry using a FACSCanto II flow cytometer (Becton Dickinson, San Jose, CA). Mitochondrial transmembrane potential was measured using tetramethylrhodamine ethyl ester (TMRE, abcam) in transient transfected HEK293T cells. Cells were incubated with 500nM of TMRE for 15min at 37°C. As a positive control for depolarizing mitochondrial membrane potential, cells were previously incubated with 20μM of FCCP (carbonilcyanide p-triflouromethoxyphenylhydrazone) for 10min at 37°C. After incubation, red fluorescence was detected in PBS/0,2% BSA by a fluorescence plate reader (Promega) at Ex/Em: 549/575nm.

**Mitochondria purification from cell lines**

Mitochondrial isolation method is based on (2). Briefly, 9 x $10^6$ HEK293T cells were culture in 150 mm plates at 80-90% confluency and transfected with different GSDMB constructs during 48h. Cells wereharvested by cell scrapper, pelleted at 600g for 5 min and washed twice in PBS. The cell pellets were placed on ice and broken by adding one volume of hypotonic homogenization buffer (IB 0.1x: 3.5 mM Tris–HCl, pH 7.8, 2.5 mM NaCl, 0.5 mM MgCl2), and homogenized by 10 strokes using a Thomas homogenizer with a motor-driven Teflon pestle. Immediately after, 1/10 of the packed cell volume of hypertonic buffer was added to make the medium isotonic. Homogenate was centrifuged at 1200g for 3 min at 4 °C to pellet unbroken cells, debris, and nuclei. Supernatant was collected and centrifuged again at low speed in the same conditions (1200g for 3 min at 4 °C). Mitochondria contained on the supernatant were pelleted in Eppendorf tubes (adding approximately 1 ml of supernatant per tube) by centrifugation in microfuge at 15 000 g during 2 min at 4°C. Pellets were washed using homogenization buffer A (0.32 M sucrose, 1 mM EDTA, and 10 mM Tris–HCl, pH 7.4) and again, resuspended in the appropriate buffer and kept at 4 ºC until used.

**Mitochondrial DNA release assay**

Human mitochondrial DNA was isolated from the cytosolic fraction of transfected HEK293T with different GSDMB constructs using the DNeasy Blood &Tissue Kit (QIAGEN). The standard of mitochondrial DNA was obtained from mitochondria isolated from HEK293T cells using the protocol described (2). Mitochondrial DNA standard curve was obtained in each assay for their absolute quantification. Quantitative PCR was employed to measure mitochondrial DNA using Power SYBR® Green PCR Master (Applied Biosystems) in a StepOnePlus™ Real-Time PCR System, using the following primers: Cytochrome c oxidase I (Forward: 5’-GCCCCAGATATAGCATTCCC-3’ and reverse: 5’-GTTCATCCTGTTCCTGCTCC-3’) and 18S rDNA (internal control) (Forward: 5’-TAGAGGGACAAGTGGCGTTC-3’ and reverse: 5’-CGCTGAGCCAGTCAGTGT-3’).

**Pyroptosis and apoptosis assays in THP1 cells**

The human non-adherent monocyte-like cell line THP1 were infected with lentiviral particles containing the following myc-tagged GSDMB plasmid constructs: 1-416, 1-416D6,7 or empty pLVX plasmid (as control). Stable expressing cells were maintained in the presence of the selection antibiotic, puromycin (12.5 µg/µL). For inducing canonical pyroptosis, $1x10^7$ cells were cultured in P100 plates with 6 mL RMPI medium in the absence of puromycin. To differentiate THP1 cells into macrophages, 0.08 µL/mL of PMA (forbol-12-miristate-13-acetate; SIGMA) were added and incubated for 24 h. Then, medium was replaced and 1 µL/mL of LPS (Lipopolysaccharide, SIGMA). After 24 hours, medium was replaced with RPMI without phenol red and containing 3 µL/mL of Nigericin (Cayman) to induce pyroptotic cell death. Control cells did not have LPS nor Nigericin. After 4 hours, medium containing dead cells was collected and centrifuged 5 minutes at 1200 rpm. The cell pellet was combined with the cells adhered to the plate, for subsequent protein extraction, as described in “protein extraction method”. The supernatant was stored at -80ºC. To quantify pyroptosis, the enzymatic activity of the released LDH (Lactate Dehydrogenase) was measured from 1 mL of the supernatant with the Cytotoxicity Detection KitPLUS (LDH) (Roche). For apoptosis induction, cells were treated with either 1 µL/mL of 10 µM etoposide (SIGMA) or DMSO, as control. After 24 hours, the supernatant containing dead cells was processed as described in the pyroptosis assays for Western blotting.

**Neutrophil Elastase protease assay**

Transient transfected cells were lysed in 0,1% Triton X-100 lysis buffer without protease inhibitors and subsequently sonicated twice during 30s (Soniprep 150). Lysates were centrifugated at 10 000 g for 10 min at 4°C and the supernatant was collected to determine the protein concentration by BCA assay (Thermo Fisher Scientific™, Carlsbad, CA, USA). Cell lysates (12,5 ug for hNE) were mixed at indicated concentrations of recombinant human neutrophile elastase (Sigma), followed by incubation at 37°C for 1h. Elastase reactions were carried out in the lysis buffer containing 0,1% Triton X-100. Where indicated, BAY-678 inhibitor was incubated with the protease treatment at the same conditions. Reactions were stopped by adding 1x Leammli Buffer with DTT (25mM) and incubated for 5 min at 95°C prior to loading on SDS-PAGE.

**Western Blot**

Total proteins were extracted in lysis buffer (0,1M NaCl, 0,05M Tris HCl pH 7,9, 5µM MgCl2, 5µM CaCl2 and 2% SDS) containing protease and phosphatase inhibitor cocktail (2mM PMSF, 2µg/ml leupeptin, 20µg/ml aprotinin, 1mM sodium orthovanadate, 5mM NaF y 5mM b-glycerophosphate; Sigma Aldrich). After 30s sonication (Soniprep 150) protein was quantified by BCA assay (Thermo Fisher Scientific™, Carlsbad, CA, USA) Membrane proteins were collected following the Plasma Membrane Protein Kit protocol (Abcam, ab65400), both, the cytosolic and membrane fractions were lysate in 0,5% Triton in PBS. Laemmli 5X buffer was finally added. Proteins were resolved in 10 or 12% SDS-PAGE and transferred on nitrocellulose membranes. After blocking the membranes with 5% skimmed milk in TBS 1X-Tween (0,01%) for 1h, primary antibodies (listed in **Supplementary Table 2)** were incubated overnight at 4°C. After washing, secondary antibodies were added at 1:3 000 for 1h at RT. Immunoblots were detected with Pierce ECL Western Blotting Substrate (Thermo Fisher Scientific™, Carlsbad, CA, USA).

**Apoptosis Caspase 3/7 detection by cytometry**

CellEvent™ Caspase 3/7 Green Flow Cytometry Assay Kit (Thermo Fisher Scientific™, Carlsbad, CA, USA) was used to detect apoptosis in HEK239T cells transfected during 48h with different GSDMB constructs. Additionally, the kit includes the SYTOX® AADvanced™ which detect death cells allowing the identification of viable cells and necrotic cells. Apoptotic, necrotic, and viable cells were measured by flow cytometry using a FACSCanto II flow cytometer (Becton Dickinson, San Jose, CA, USA).

**Immunofluorescence, confocal microscopy and Cell Observer**

Cells were seeded in 24-well cell culture plate (1,5 x $10^5$ cell/well) andtransfected for 72 hours. Then cells were washed with PBS and incubated with red MitoTracker™ Deep Red FM (Thermo Fisher Scientific™, Carlsbad, CA, USA) for 30 min at 37°C. Next, cells were fixed with 4% paraformaldehyde 20 min at room temperature. After 15 min of permeabilization with Triton X100 0,1%, cells were washed with PBS and incubated with primary antibodies (detailed in **Supplementary Table 2)** for 90 min at room temperature. Then, secondary antibodies were incubated for 45 min at 1:1 000. Immunofluorescence preparations were visualized in a confocal microscopy LSM710 (Zeiss) and images were processed by Fiji software (Image J 1.52).

To assess GSDMB intracellular localization in real time, HEK293T cells were transiently transfected with Doxycycline-inducible vectors expressing GFP-tagged GSDMB constructs. Cells were pre-induced with Doxycycline at 200ng for 3h. Additionally, to assess the dynamics of cell death induction, cells were cultured in the presence of 0.2µg/ml Propidium Iodide. Live videos (1 frame every 10 minutes for at least 20h) were recorded with either Cell Observer Microscopy (ZEISS, Oberkochen Germany) or LSM710 confocal microscopy (ZEISS). Videos were processed by Microscope Software ZEN lite (ZEISS) and Fiji software (Image J 1.52), respectively.

**Correlative light and electron microscopy (CLEM)**

For correlative light and electron microscopy (CLEM) studies in 23132/87 cells were seeded in a permanox Lab-Tek chamber slide of 4 wells (Nalge Nunc International, Naperville, IL) at a density of 1,5 x $10^5$ cell/well and transfected with doxycycline inducible vectors 1-220-GFP and 1-242-GFP as previously mentioned. After 6h of transfection, cells were induced with Doxycycline at 200ng/ml. Cells were incubated with red MitoTracker™ Deep Red FM (Thermo Fisher Scientific™, Carlsbad, CA, USA) for 30 min at 37°C. Finally, the cells were fixed in 3 % glutaraldehyde in 0.1 M phosphate buffer (PB) for 1 hour at 37ºC. Last, were washed in 0.1 M phosphate buffer (PB) for 5 times and stored at 4 ºC.

The slides were assembled with Dako Mounting Medium containing DAPI. The confocal images were acquired with a Leica TCS SP8 HyVolution II (Leica Microsystems, Wetzlar, Germany) inverted laser scanning confocal microscope using oil objective 63X Plan-Apochromat-Lambda Blue 1.4 N.A. The excitation wavelengths for fluorochromes were 488 nm for GFP, 638 nm for Mitotracker Deep Red and 405 nm for Dapi. Optical sections were acquired every 0.346 µm. Two-dimensional pseudo color images (255 color levels) were gathered with a size of 1024x1024 pixels, 2X optical zoom and Airy 1 pinhole diameter. After fluorescence capture, slides were processed for transmission electron microscopy analysis. The samples were postfixed in 2% OsO4 for 1h at room temperature and stained in 2% uranyl acetate in the dark for 2h at 4 °C. Then, were rinsed in distilled water, dehydrated in ethanol, and infiltrated overnight in Durcupan resin (Sigma-Aldrich, St. Louis, USA). Following polymerization, embedded cultures were detached from the wells and glued to Durcupan blocks. Finally, ultrathin sections (0.08 µm) were cut with an Ultracut UC-6 (Leica microsystems, Wetzlar, Germany), stained with lead citrate (Reynolds solution) and examined under a transmission electron microscope FEI Tecnai Spirit BioTwin (Thermo Fisher Scientific™, Carlsbad, CA, USA). Pictures were taken using Radius software (Version 2.1) with a Xarosa digital camera (EMSIS GmbH, Münster, Germany).

**qRT-PCR in human samples**

Gene expression was analyzed by real-time PCR, using StepOnePlus Real-time PCR System, with TaqMan® Fast Advanced Master Mix (Applied Biosystems™ by Thermo Fisher Scientific™, Carlsbad, CA, USA). Normalization was done with *GAPDH*. Relative expression was calculated using the comparative Ct method and obtaining the fold-change value (ΔΔCt). TaqMan probes used for qPCR assessment of *GSDMB* expression and *GSDMB* isoforms are listed in **Supplementary Table 2**.

**In-Gel Digestion and Reverse phase-liquid chromatography RP-LC-MS/MS analysis**

After drying, gel bands or spots were distained in acetonitrile:water (ACN:H2O, 1:1), were reduced and alkylated (disulphide bonds from cysteinyl residues were reduced with 10 mM DTT for 30 min at 56 ºC, and then thiol groups were alkylated with 10 mM iodoacetamide for 30 min at room temperature in darkness) and digested in situ with sequencing grade trypsin (Promega, Madison, WI) as described by (3) with minor modifications (4). The gel pieces were shrunk by removing all liquid using sufficient ACN. Acetonitrile was pipetted out and the gel pieces were dried in a speedvac. The dried gel pieces were re-swollen in 100 mM Tris-HCl pH 8, 10mM $CaCl_2$ with 12.5 ng/µl trypsin for 1h in an ice-bath. The digestion buffer was removed, and gels were covered again with 100 mM Tris-HCl pH 8, 10mM $CaCl_2$ and incubated for 12 h at 37°C. Digestion was stopped by the addition of 1% TFA. Whole supernatants were dried down and then desalted onto ZipTip C18 Pipette tips (Millipore) until the mass spectrometric analysis. The desalted protein digest was dried, resuspended in 10 µl of 0.1% formic acid and analyzed by RP-LC-MS/MS in an Easy-nLC II system coupled to an ion trap LTQ-Orbitrap-Velos-Pro hybrid mass spectrometer (Thermo Fisher Scientific™, Carlsbad, CA, USA). The peptides were concentrated (on-line) by reverse phase chromatography using a 0.1mm × 20 mm C18 RP precolumn (Thermo Fisher Scientific™, Carlsbad, CA, USA), and then separated using a 0.075mm x 250 mm C18 RP column (Thermo Fisher Scientific™, Carlsbad, CA, USA) operating at 0.3 µl/min. Peptides were eluted using a 60-min dual gradient. The gradient profile was set as follows: 5−25% solvent B for 68 min, 25−40% solvent B for 22 min, 40−100% solvent B for

2min and 100% solvent B for 18 min (Solvent A: 0,1% formic acid in water, solvent B: 0,1% formic acid, 80% acetonitrile in water). ESI ionization was done using a Nano-bore emitters Stainless Steel ID 30 µm (Proxeon) interface at 2.1 kV spray voltage with S-Lens of 60%. The Orbitrap resolution was set at 30 000 (5).

**Breast cancer TCGA data analysis**

GSDMB isoforms mRNA expression from breast cancer patients (N=1 093) was analyzed using data from TCGA. Correlation of GSDMB isoforms and overall survival was analyzed using "survival" R package (R Bioconductor). P-values were calculated using long-rank test. Hazard rates were calculated by Cox proportional-hazards model. GSDMB isoforms expression was scored according to the mean expression. Results were considered significant when $p\text{-value} < 0.05$.

**Supplementary Figures:**

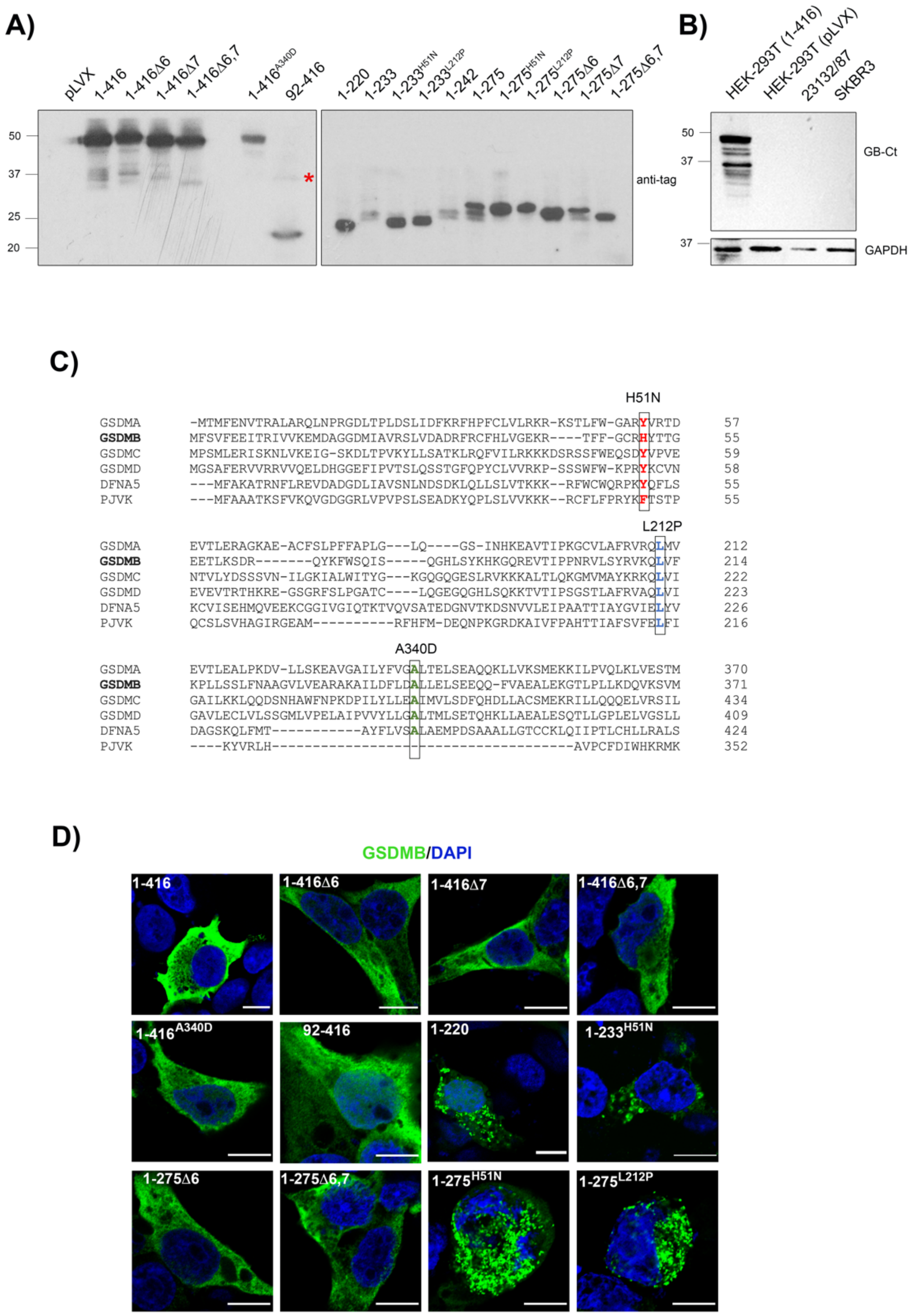


**Supplementary Figure 1. A**) Western blot representation of all constructs included in the study. GSDMB constructs were transiently transfected during 48 h in HEK293T cell lines. Protein expression was analyzed by Western Blot using anti-GSDMB detecting CT region antibody DAN

114B (6) or the anti-myc or anti-HA antibody detecting the tag from the CT. *Band expected for the construct 92-416. **B**) Western blot showing GSDMB endogenous protein expression in the different cell lines. HEK293T transiently transfected with GSDMB-full length (1-416) was used as a positive control. GSDMB was detected using anti-GSDMB-CT antibody DAN 114B (6). **C**) Sequence alignment of the six GSDMs members. Squares indicate point mutation position in all GSDMs. **D**) Subcellular localization of GSDMB constructs in HEK293T cells transiently transfected during 48h. Localization was analyzed by immunofluorescence and confocal microscopy. GSDMB NT (green) was detected with SIGMA antibody (HPA023925). Nuclei (blue) was stained with DAPI. Scale bar represents 10μm.

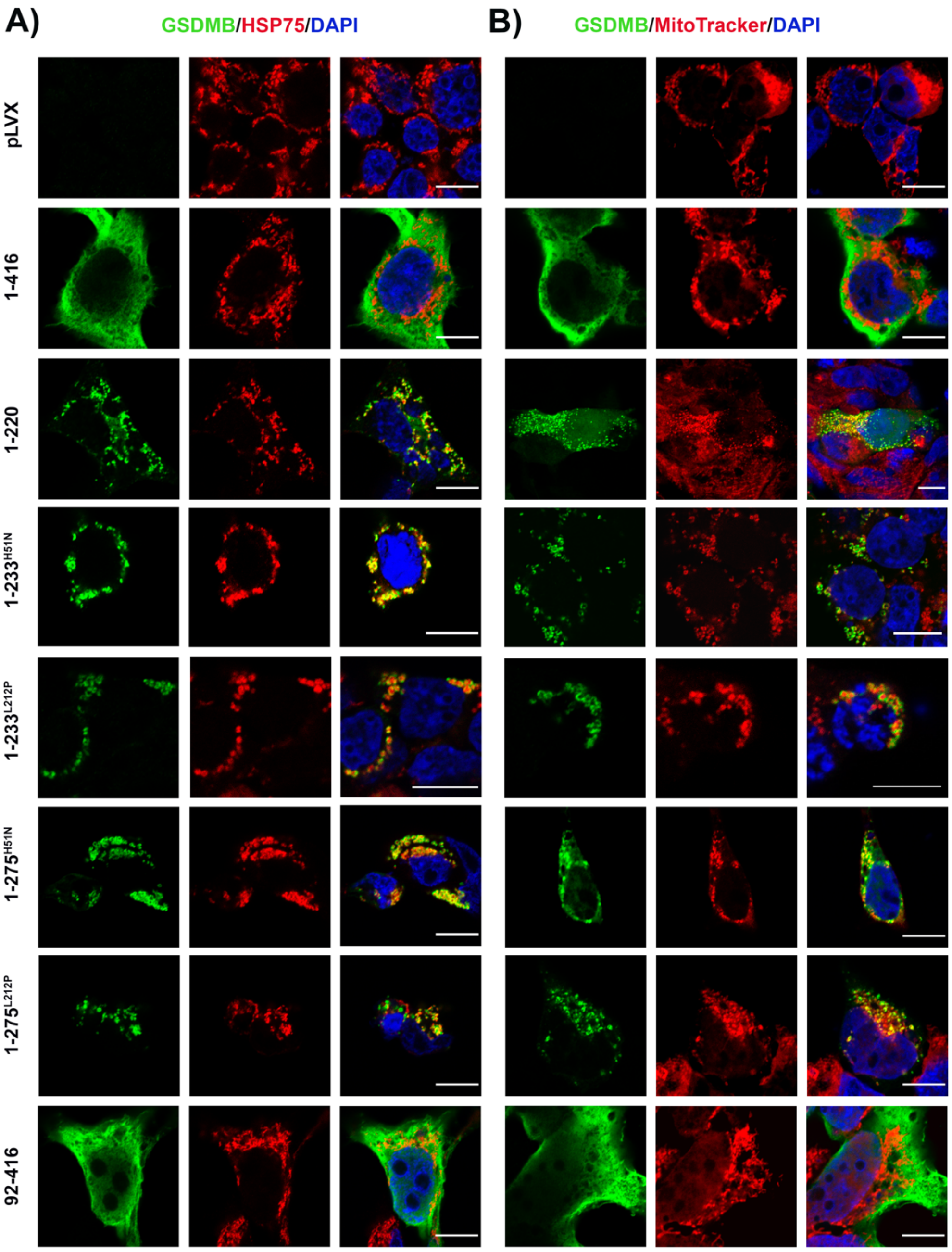


**Supplementary Figure 2. GSDMB-NT aggregates co-localize with mitochondrial markers Hsp75/Trap1 and MitoTracker.** HEK293T cells were transiently transfected (48h) with the indicated GSDMB constructs. Intracellular localization of GSDMB constructs (green; NT antibody SIGMA, HPA023925) and co-localization with mitochondria (red) Hsp75 (**A**) and MitoTracker Deep Red (**B**) by immunofluorescence and confocal microscopy analysis. Cells transfected with empty vector (pLVX) were used as a negative control. GSDMB-Full length protein (1-416) and CT region (92-416) do not co-localize with mitochondrial markers. Nuclei (blue) was stained with DAPI. Scale bar represents 10μm.

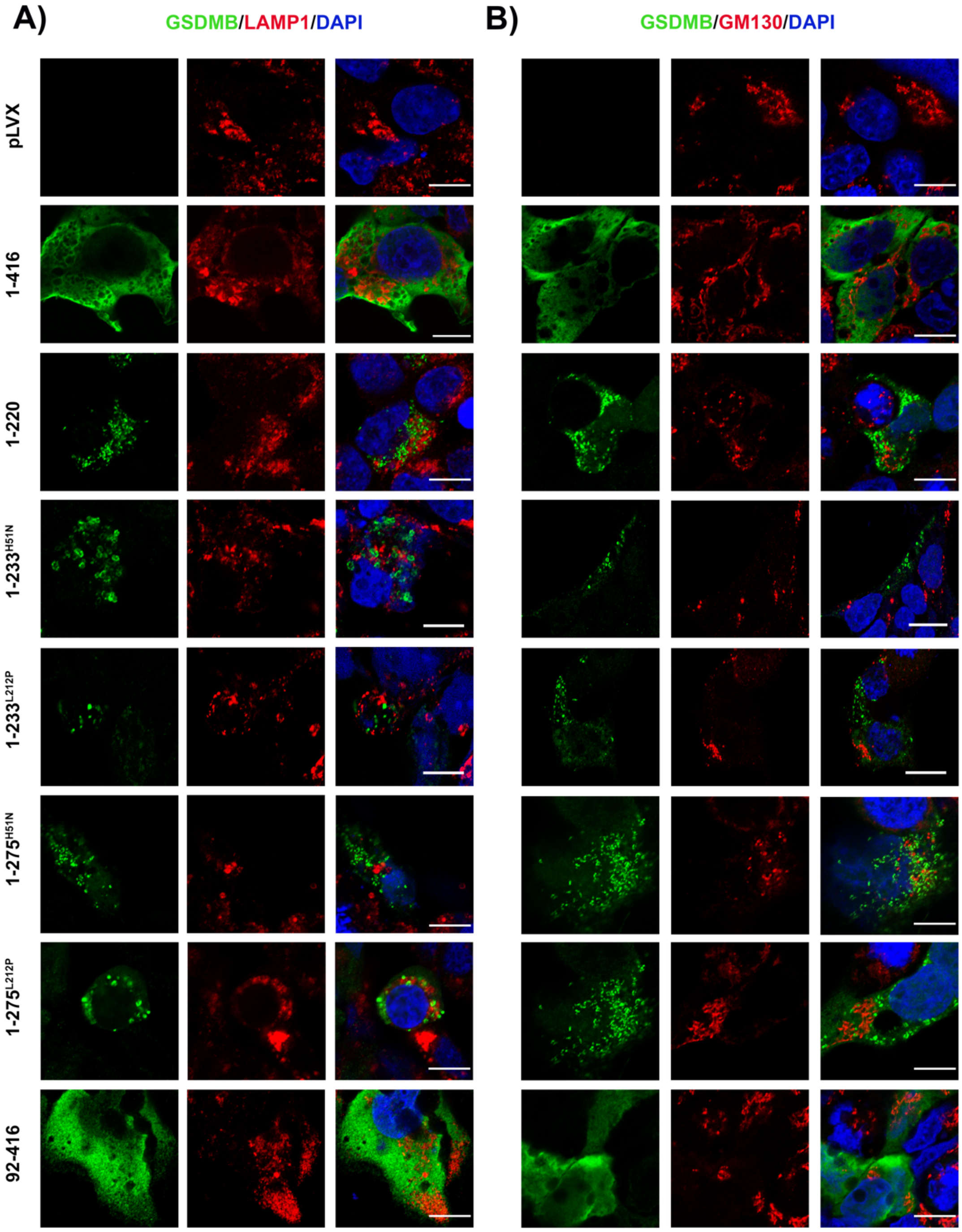


**Supplementary Figure 3. GSDMB-NT aggregates do not co-localize with lysosome (LAMP1) or Golgi (GM130) markers.** HEK293T cells were transiently transfected (48h) with the indicated GSDMB constructs. Intracellular localization of GSDMB constructs (green; NT antibody SIGMA, HPA023925), lysosomes (red, LAMP) (**A**) and Golgi (red, GM130) (**B**) by immunofluorescence and confocal microscopy analysis. Nuclei (blue) was stained with DAPI. Scale bar represents 10μm.

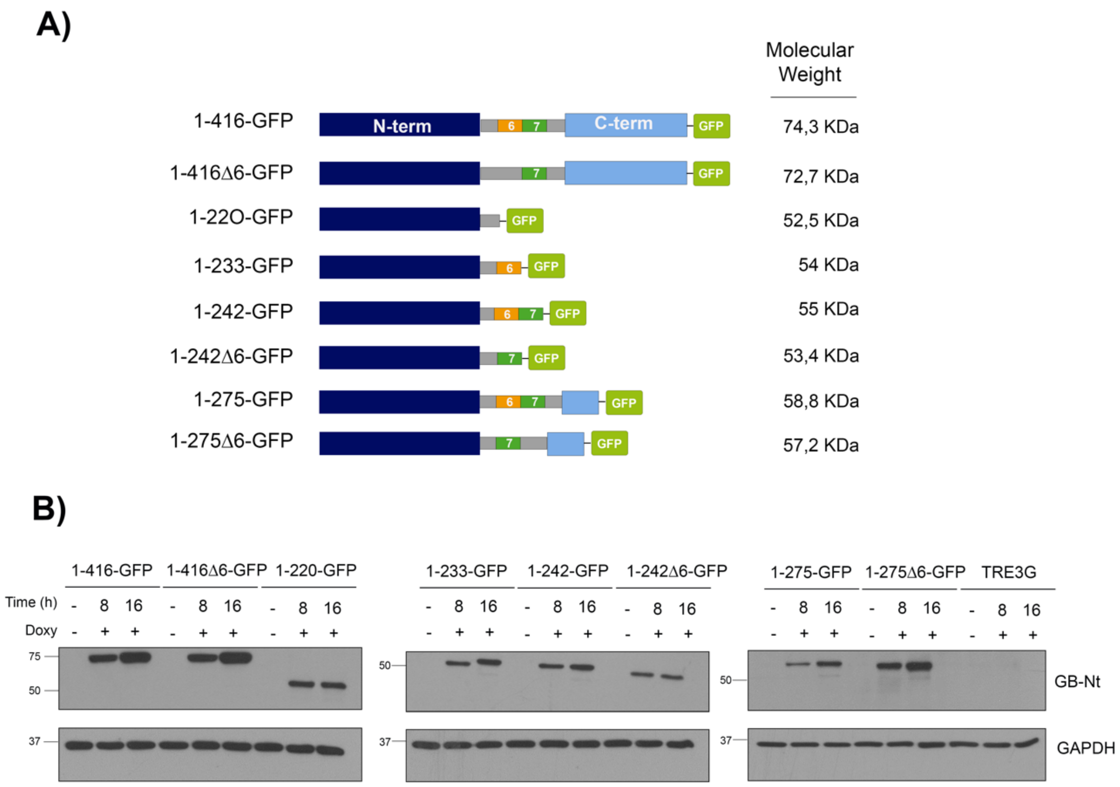


**Supplementary Figure 4. Doxycycline inducible GSDMB constructs fused (in the CT) with green fluorescence protein (GFP)**. **A**) Schematic representation of the constructs. **B**) Western-blot confirming the inducible expression of GSDMB-GFP constructs. HEK293T were transiently transfected (24h) and GSDMB-GFP was induced by Doxycycline at 200ng/ml during 8 or 16h. GSDMB-GFP constructs were detected using anti-GSDMB detecting NT region (Santa Cruz Biotech, sc-101239). GAPDH was used as a loading control.

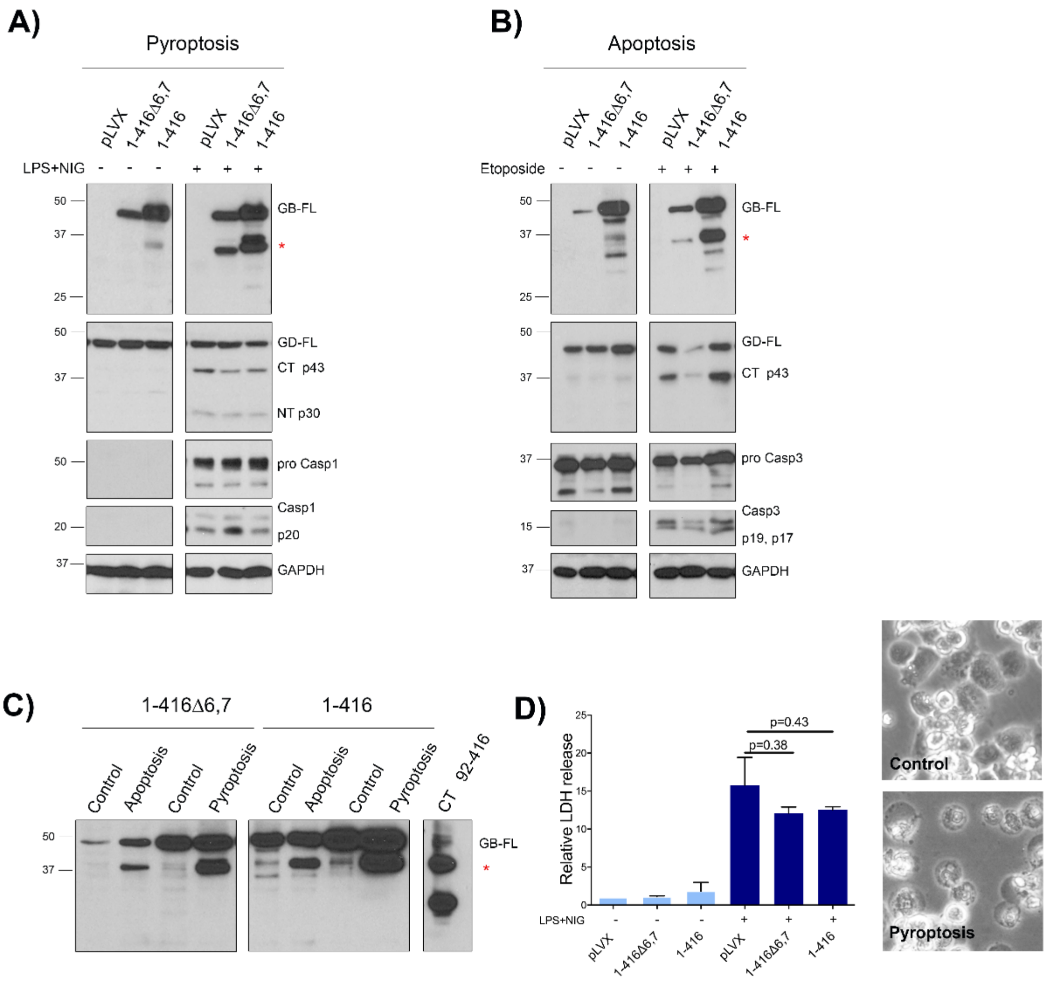


**Supplementary Figure 5. GSDMB is cleaved in a similar way during pyroptosis and apoptosis and its over-expression does not affect the extent of pyroptosis in THP1 cells.**
**A-B)** THP1 stably expressing either GSDMB isoform 2 (1-275Δ6,7), isoform 3 (1-416) or the empty vector (pLVX) were induced to undergo pyroptosis (**A:** 24h LPS and 4h Nigericin) or apoptosis (**B:** 10μM etoposide or DMSO for 24h) and cell lysates were subject to Western Blot. Using an anti-GSDMB-CT antibody DAN 114B (6) a CT fragment (asterisk) of 35 KDa (1-275D6,7) and 37 KDa (1-416) was detected in both pyroptosis or apoptosis assays. For comparison, GSDMD showed a differential cleavage pattern between apoptosis (p43 CT fragment) and pyroptosis (p30 NT fragment and secondary p43 CT fragment), as reported before (7). Pro-caspase 1 and activated caspase-1 were detected in the supernatant from pyroptotic cells (**A**). Caspase 3 activation was detected in cell lysates undergoing apoptosis (**B**). GAPDH was used as a loading control. Molecular weights are shown on the left. **C)** Comparison in parallel of

cell lysates from the pyroptosis and apoptosis experiments shown in (**A-B**) for each GSDMB isoform, confirming that the detected CT fragment (asterisk) equals to the 92-416 construct (which corresponds to the CT peptide generated by caspases 1/3/4/7/8/9 (8, 9). **D)** No significant effect of GSDMB isoform over-expression on the extent of pyroptotic cell lysis measured by LDH release. Canonical pyroptosis was induced by LPS and Nigericin treatment. Control cells were grown under standard culture conditions. Bars represents mean values ± SD from three independent experiments. Differences between control (pLVX) and GSDMB-expressing conditions were tested by t-test. Phase contrast photographs taken of control treated cells (top) and LPS+Nigericin treated cells (bottom). Note the changes on morphology and adhesion, typical of pyroptosis.

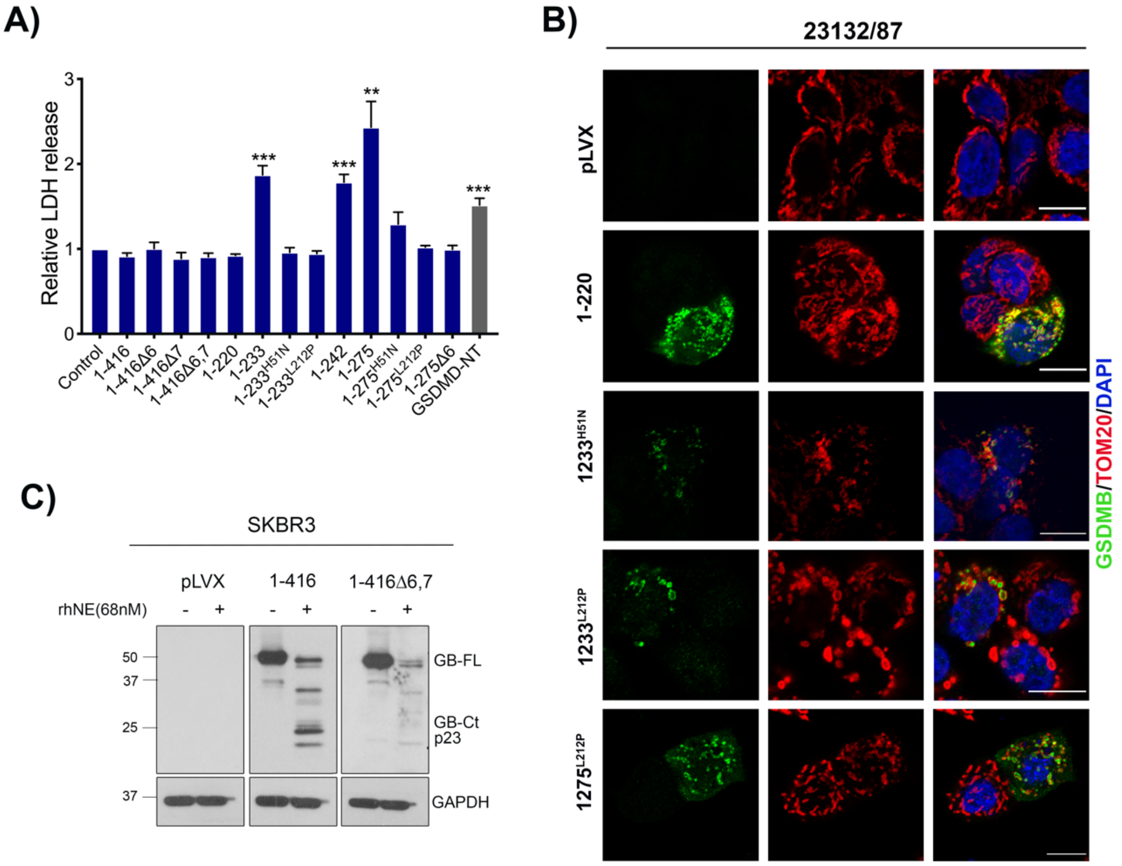


**Supplementary Figure 6. Pro-cell death role of GSDMB-NT in cancer cells in an isoform-dependent way.** Gastric cancer cell line 23132/87 was transiently transfected with GSDMB constructs during 48h. **A**) Cytotoxicity was measured by lactate dehydrogenase (LDH) assay. Bars represents mean values ± SEM from more than three independent experiments. Differences between control condition (empty vector) and each condition was tested by t-test: **p<0,01 and ***p<0,001. GSDMD-NT was used as a positive control. **B**) Immunofluorescence and confocal microscopy analysis in 23132/87 cells transiently transfected with indicated GSDMB-NT constructs. GSDMB-NT (green; NT antibody SIGMA, HPA023925), co-localizes with mitochondrial marker TOM20 in red. pLVX was the empty vector used as a negative control. **C**) Immunoblotting analysis of GSDMB cleavage by recombinant human neutrophil elastase (rhNE). SKBR3 breast cancer cell lysates containing GSDMB isoforms 2 and 3 (without exon 6 and with exon 6, respectively) were incubated with rhNE (68nM) at 37°C for 30 min. GSDMB-CT p23 fragment generated was detected using anti-GSDMB-CT antibody DAN 114B (6).

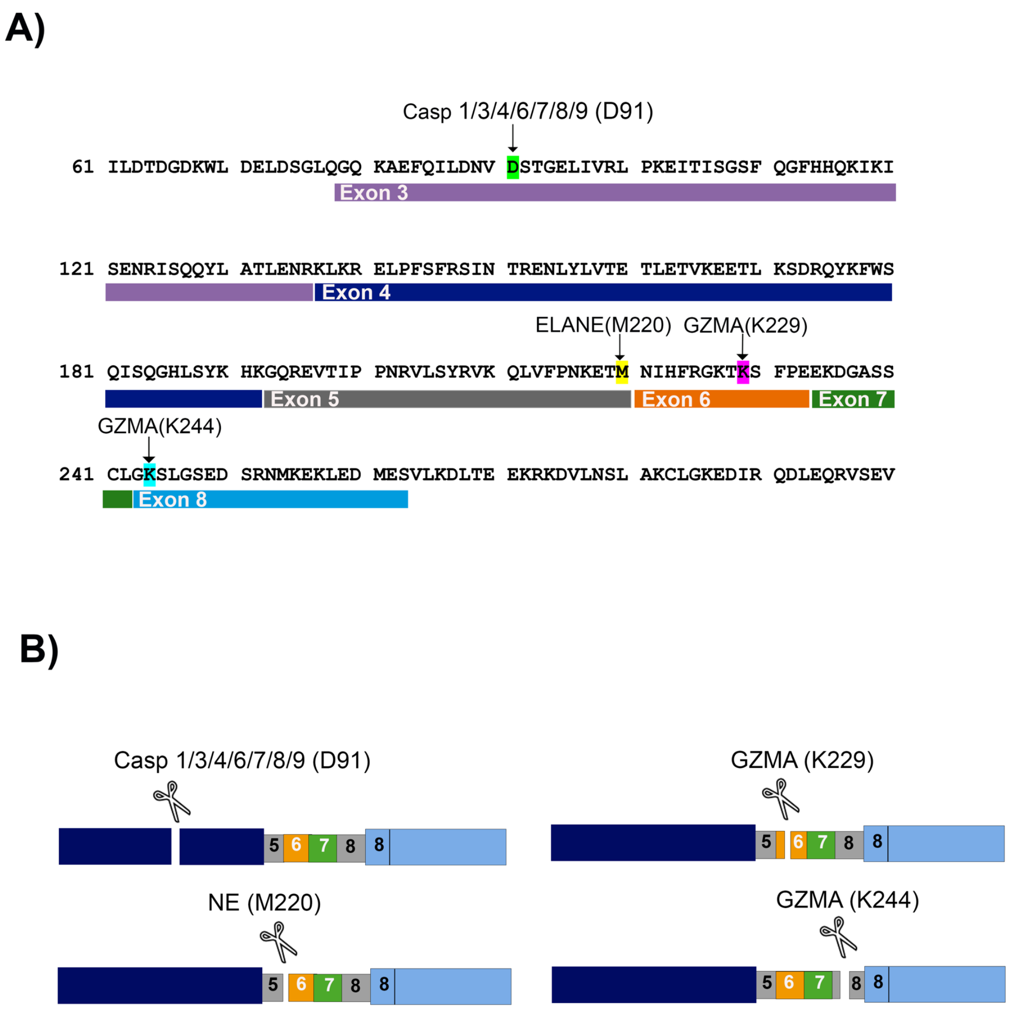


**Supplementary Figure 7.** GSDMB cleavage by GZMA, Neutrophil Elastase (NE) and caspases. **A**) GSDMB sequence and cleavage sites representation indicating the exon localization. **B**) Schematic representation of GSDMB cleavage by GZMA, NE and caspases.

### Supplementary Tables:

**Supplementary Table 1:** Summary table of functional activity of GSDMB constructs and subcellular localization.

| GSDMB constructions | Cell localization (HEK293T) | Citotoxicity (LDH) | Mitochondrial damage (mitoSOX) | Membrane Potential (TMRE) |
|---|---|---|---|---|
| 1-416 | Diffuse cytoplasmic | No | No | No |
| 1-416 Δ6 | Diffuse cytoplasmic | No | No | No |
| 1-416 Δ7 | Diffuse cytoplasmic | No | No | No |
| 1-416 Δ6,7 | Diffuse cytoplasmic | No | No | No |
| $1\text{-}416^{A340D}$ | Diffuse cytoplasmic | No | No | - |
| 92-416 | Diffuse cytoplasmic | No | No | - |
| 1-220 | Mitochondrial aggregates | No | No | No |
| 1-233 | No detected | High* | High* | Decreased* |
| $1\text{-}233^{H51N}$ | Mitochondrial aggregates | No | No | No |
| $1\text{-}233^{L212P}$ | Mitochondrial aggregates | No | No | No |
| 1-242 | No detected | High* | High* | Decreased* |
| 1-275 | No detected | High* | High* | Decreased* |
| $1\text{-}275^{H51N}$ | Mitochondrial aggregates | No | No | No |
| $1\text{-}275^{L212P}$ | Mitochondrial aggregates | Moderate* | No | No |
| 1-275, Δ6 | Diffuse cytoplasmic | No | No | No |
| 1-275 Δ7 | No detected | High* | High* | No |
| 1-275 Δ6,7 | Diffuse cytoplasmic | No | No | No |

*Results with significant p-value<0.05

**Supplementary Table 2:** Key resources table.

| Reagent or resource | Source | Identifier | Probe and dilution |
|---|---|---|---|
| Antibodies | | | |
| anti Caspase 3 | Santa Cruz Biotech | sc-56053 | WB 1:250 |
| anti GAPDH | Calbiochem | CB1001 | WB 1:50 000 |
| anti Gasdermin B | in house | DAN/114B (7) | WB 1:250 |
| anti Gasdermin B | Abcam | ab215729 | WB 1:1 000 |
| anti Gasdermin B | Santa Cruz Biotech | sc-101239 | WB 1:250 |
| anti Gasdermin B | Sigma Aldrich | HPA023925 | IF 1:50 |
| anti GFP | Invitrogen | A11122 | WB 1:2 000 |
| anti GM130 | BD BIOSCIENCES | 610822 | IF 1:100 |
| anti Granzyme A | Abcam | ab209205 | WB 1:1 000 |
| anti HA | Sigma Aldrich | 12158167001 | WB 1:500 |
| anti Hsp90a/b | Santa Cruz Biotech | sc-13119 | WB 1:500 |
| anti human Neutrophile Elastase | Santa Cruz Biotech | sc-55549 | WB 1:500 |
| anti LAMP1 | Santa Cruz Biotech | sc-20011 | IF 1:200 |
| anti myc Tag (mouse) | Cell Signaling | 2276 | WB 1:1 000 |
| anti myc Tag (rabbit) | Cell Signaling | 2278S | WB 1:1 000 |
| anti Trap1/Hsp75 | Santa Cruz Biotech | sc-390061 | IF 1:100/WB 1:50 |
| anti TOM20 | Santa Cruz Biotech | sc-17764 | IF 1:200/WB 1:200 |
| anti α1 Sodium Potassium ATPase | Abcam | ab7671 | WB 1:10 000 |
| anit α-Tubulin | Sigma Aldrich | T9026 | WB 1:10 000 |
| Mouse IgGκ light chain | Santa Cruz Biotech | sc-516102 | WB 1:3 000 |
| Mouse anti-rabbit | Santa Cruz Biotech | sc-2357 | WB 1:3 000 |
| Goat anti-rat | Nordic Mubio | GARa/IgG2a/PO | WB 1:3 000 |
| Cell lines | | | |
| HEK293T | ATCC- American Type Culture Collection | ATCC CRL-3216 | |
| SKBR3 | DSMZ-German Collection of Microorganisms | ACC 736 | |
| 23132/87 | DSMZ-German Collection of Microorganisms | ACC 201 | |
| NK-92 | DSMZ-German Collection of Microorganisms | ACC 488 | |
| THP1 | ATCC- American Type Culture Collection | ATCC TIB-202 | |
| Plasmids | | | |
| pEZ-M61 | Genecopoeia | EX-NEG-M61 | |
| pLVX-Puro-MYC | Clontech, Takara | 632164 | |
| Lenti-TRE3G-GFP | Transomic | TLO3092-TRE3G-ORF-C-TagGFP2-PGK-Tet3G-puro | |
| pCS2-3XFlag-hGSDMD | Dr. Pablo Pelegrin | - | |
| pCMV6-XL4 | Origene | NC1720451 | |
| qRT PCR reagents and probes | | | |
| GSDMB ISOFORM1 | Applied Biosystems™ | Hs00938445_m1 | |
| GSDMB ISOFORM2 | Applied Biosystems™ | Hs00939390_m1 | |
| GSDMB ISOFORM3 and 4 | Applied Biosystems™ | Hs00940508_m1 | |
| GAPDH | Applied Biosystems™ | Hs02758991_g1 | |
| TaqMan® Fast Advanced Master Mi: | Applied Biosystems™ | 4444556 | |

**Supplementary Videos legends:**

**Supplementary Video 1.** Live-cell imaging of HEK293T cells expressing 1-416-GFP after 2 hours of pre-induction with doxycycline. Cells were recorded for at least 20h (1 picture every 10-30 minutes). Propidium Iodide (0.2µg/ml in red) stains nuclei from dying cells. Bright field shown. Scale bar represents 10µm.

**Supplementary Video 2**. Live-cell imaging of HEK293T cells expressing 1-416-GFP after 2 hours of pre-induction with doxycycline using Cell Observer system for 1-2 hours. Propidium Iodide (0.2µg/ml in red) stains nuclei from dying cells. Scale bar represents 10µm.

**Supplementary Video 3.** Live-cell imaging of HEK293T cells expressing 1-220-GFP after 2 hours of pre-induction with doxycycline using Cell Observer system for 1-2 hours. Propidium Iodide (0.2µg/ml in red) stains nuclei from dying cells. Bright field shown. Scale bar represents 10µm.

**Supplementary Video 4**. Live-cell imaging of HEK293T cells expressing 1-220-GFP after 2 hours of pre-induction with doxycycline using Cell Observer system for 1-2 hours. Propidium Iodide (0.2µg/ml in red) stains nuclei from dying cells. Scale bar represents 10µm.

**Supplementary Video 5.** Live-cell imaging of HEK293T cells expressing 1-242Δ6-GFP after 2 hours of pre-induction with doxycycline using Cell Observer system for 1-2 hours. Propidium Iodide (0.2µg/ml in red) stains nuclei from dying cells. Bright field shown. Scale bar represents 10µm.

**Supplementary Video 6**. Live-cell imaging of HEK293T cells expressing 1-242Δ6-GFP after after 2 hours of pre-induction with doxycycline using Cell Observer system for 1-2 hours. Propidium Iodide (0.2µg/ml in red) stains nuclei from dying cells. Scale bar represents 10µm.

**Supplementary Video 7.** Live-cell imaging of HEK293T cells expressing 1-275Δ6-GFP after 2 hours of pre-induction with doxycycline using Cell Observer system for 1-2 hours. Propidium Iodide (0.2µg/ml in red) stains nuclei from dying cells. Bright field shown. Scale bar represents 10µm.

**Supplementary Video 8**. Live-cell imaging of HEK293T cells expressing 1-275Δ6-GFP after 2 hours of pre-induction with doxycycline using Cell Observer system for 1-2 hours. Propidium Iodide (0.2µg/ml in red) stains nuclei from dying cells. Scale bar represents 10µm.

**Supplementary Video 9**. Live-cell confocal imaging over 1-2 hours of HEK293T cells expressing 1-242-GFP following doxycycline induction. Propidium Iodide (0.2µg/ml in red) stains nuclei from dying cells. Bright field shown. Scale bar represents 10µm.

**Supplementary Video 10**. Live-cell confocal imaging over 1-2 hours of HEK293T cells expressing 1-242-GFP following doxycycline induction. Propidium Iodide (0.2µg/ml in red) stains nuclei from dying cells. Scale bar represents 10µm.

**Supplementary Video 11**. Live-cell confocal imaging over 1-2 hours of HEK293T cells expressing 1-275-GFP following doxycycline induction. Propidium Iodide (0.2µg/ml in red) stains nuclei from dying cells. Bright field shown. Scale bar represents 10µm.

**Supplementary Video 12**. Live-cell confocal imaging over 1-2 hours of HEK293T cells expressing 1-275-GFP following doxycycline induction. Propidium Iodide (0.2μg/ml in red) stains nuclei from dying cells. Scale bar represents 10μm.
